# Alternative hydrogen uptake pathways suppress methane production in ruminants

**DOI:** 10.1101/486894

**Authors:** Chris Greening, Renae Geier, Cecilia Wang, Laura C. Woods, Sergio E. Morales, Michael J. McDonald, Rowena Rushton-Green, Xochitl C. Morgan, Satoshi Koike, Sinead C. Leahy, William J. Kelly, Isaac Cann, Graeme T. Attwood, Gregory M. Cook, Roderick I. Mackie

## Abstract

Farmed ruminants are the largest source of anthropogenic methane emissions globally. The methanogenic archaea responsible for these emissions use molecular hydrogen (H_2_), produced during bacterial and eukaryotic carbohydrate fermentation, as their primary energy source. In this work, we used comparative genomic, metatranscriptomic, and co-culture-based approaches to gain a system-wide understanding of the organisms and pathways responsible for ruminal H_2_ metabolism. Two thirds of sequenced rumen bacterial and archaeal genomes encode enzymes that catalyze H_2_ production or consumption, including 26 distinct hydrogenase subgroups. Metatranscriptomic analysis confirmed that these hydrogenases are differentially expressed in sheep rumen. Electron-bifurcating [FeFe]-hydrogenases from carbohydrate-fermenting Clostridia (e.g. *Ruminococcus*) accounted for half of all hydrogenase transcripts. Various H_2_ uptake pathways were also expressed, including methanogenesis (*Methanobrevibacter*), fumarate reduction and nitrate ammonification (*Selenomonas*), and acetogenesis (*Blautia*). Whereas methanogenesis predominated in high methane yield sheep, alternative uptake pathways were significantly upregulated in low methane yield sheep. Complementing these findings, we observed significant differential expression and activity of the hydrogenases of the hydrogenogenic cellulose fermenter *Ruminococcus albus* and the hydrogenotrophic fumarate reducer *Wolinella succinogenes* in co-culture compared to pure culture. We conclude that H_2_ metabolism is a more complex and widespread trait among rumen microorganisms than previously recognized. There is evidence that alternative hydrogenotrophs, including acetogens and selenomonads, can prosper in the rumen and effectively compete with methanogens for H_2_ in low methane yield ruminants. Strategies to increase flux through alternative H_2_ uptake pathways, including animal selection, dietary supplementation, and methanogenesis inhibitors, may lead to sustained methane mitigation.

## Introduction

Methane production by livestock accounts for over 5% of global greenhouse gas emissions annually ^1^. These emissions mostly originate from the activity of methanogens within ruminants, which generate methane as an obligate end-product of their energy metabolism ^2^. Several lineages of methanogenic archaea are core members of the microbiome of the ruminant foregut ^3–5^. Of these, hydrogenotrophic methanogens are dominant in terms of both methane emissions and community composition ^6,7^, with global surveys indicating that *Methanobrevibacter gottschalkii* and *Methanobrevibacter ruminantium* comprise 74% of the rumen methanogen community ^5^. These organisms use molecular hydrogen (H_2_) to reduce carbon dioxide (CO_2_) to methane through the Wolfe cycle of methanogenesis ^8,9^. Rumen methanogens have also been identified that use formate, acetate, methyl compounds, and ethanol as substrates, but usually do so in conjunction with H_2_ ^5,10–12^. Given their major contribution to greenhouse gas emissions, multiple programs are underway to mitigate ruminant methane production ^13,14^. To date, most strategies have focused on direct inhibition of methanogens using chemical compounds or vaccines ^15–18^. A promising alternative strategy is to modulate the supply of substrates to methanogens, such as H_2_, for example through dietary or probiotic interventions ^14,19,20^. To achieve this, while maintaining productivity of the host animal, requires an understanding of the processes that mediate substrate supply to methanogens within the rumen.

H_2_, the main substrate supporting ruminal methanogenesis, is primarily produced through fermentation processes ^6^. Various carbohydrate fermentation pathways lead to the production of H_2_ as an end-product, together with volatile fatty acids (VFAs) and CO_2_ ^21–23^. This process is supported by hydrogenases, which reoxidize cofactors reduced during carbohydrate fermentation and dispose of the derived electrons by producing H_2_. While it is unclear which rumen microorganisms mediate H_2_ production *in situ*, a range of isolates have been shown to produce H_2_ *in vitro* ^24–28^. For example, the model rumen bacterium *Ruminococcus albus* 7 reoxidizes the reduced ferredoxin and NADH formed during glucose fermentation using two different [FeFe]-hydrogenases depending on environmental conditions ^29^. In addition, it is well-established that some rumen fungi and ciliates produce H_2_ *via* hydrogenosomes ^30,31^.

A further potential source is the nitrogenase reaction, which produces one H_2_ for every N_2_ fixed; however, while numerous rumen microorganisms encode putative nitrogenases ^21^, there is no convincing *in situ* evidence that N_2_ fixation occurs in the rumen ^32^. A large proportion of the H_2_ produced by hydrogenogenic fermenters is directly transferred to hydrogenotrophic methanogens, in an ecological process known as interspecies hydrogen transfer ^25,33^. Particularly remarkable are the endosymbiotic and ectosymbiotic associations of methanogens, such as *M. ruminantium*, with rumen ciliates ^34–36^. In addition to providing a continual substrate supply for methanogens, such symbioses benefit fermenters by maintaining H_2_ at sufficiently low concentrations for fermentation to remain thermodynamically favorable ^37^.

Various hydrogenotrophic bacteria are thought to compete with methanogens for the rumen H_2_ supply. Most attention has focused on homoacetogens, which mediate conversion of H_2_/CO_2_ to acetate using [FeFe]-hydrogenases ^38^. Several genera of homoacetogens have been isolated from the rumen, including *Eubacterium* ^39^, *Blautia* ^40^, and *Acetitomaculum* ^41^. However, molecular surveys indicate their abundance is generally lower than hydrogenotrophic methanogens ^42–44^. This is thought to reflect that methanogens outcompete acetogens due to the higher free energy yield of their metabolic processes, as well as their higher affinity for H_2_. The dissolved H_2_ concentration fluctuates in the rumen depending on diet, time of feeding, and rumen turnover rates, but is generally at concentrations between 400 to 3400 nM ^45^; these concentrations are typically always above the threshold concentrations required for methanogens (< 75 nM) but often below those of homoacetogens (< 700 nM) ^46^. Despite this, it has been proposed that stimulation of homoacetogens may be an effective strategy for methane mitigation in methanogen-inhibited scenarios ^14,20,47,48^. Various microorganisms have also been isolated from cows and sheep that support anaerobic hydrogenotrophic respiration, including dissimilatory sulfate reduction (e.g. *Desulfovibrio desulfuricans*) ^49,50^, fumarate reduction and nitrate ammonification (e.g. *Selenomonas ruminantium, Wolinella succinogenes*) ^51–58^, and trimethylamine *N*-oxide reduction (e.g. *Denitrobacterium detoxificans*) ^59^. The first described and most comprehensively studied of these hydrogen oxidizers is *W. succinogenes*, which mediates interspecies hydrogen transfer with *R. albus* ^25^. In all cases, respiratory electron transfer *via* membrane-bound [NiFe]-hydrogenases and terminal reductases generates a proton-motive force that supports oxidative phosphorylation ^60^. It is generally assumed that these pathways are minor ones and are limited by the availability of oxidants. Promisingly, it has been observed that dietary supplementation with fumarate, sulfate, or nitrate can significantly reduce methane production in cattle, likely by stimulating alternative pathways of H_2_ consumption ^61,62^.

We postulate that mitigating methane emissions, while maintaining animal productivity, depends on understanding and controlling H_2_ utilization by methanogens. This requires a system-wide perspective of the schemes for production and concomitant utilization of H_2_ in the rumen. To facilitate this, we determined which organisms and enzymes are primarily responsible for H_2_ production and consumption in rumen. Firstly, we screened genome, metagenome, and metatranscriptome datasets ^21,63,64^ to resolve the microbial genera, metabolic pathways, and hydrogenase classes ^65,66^ that mediate H_2_ metabolism. We demonstrate that ruminants harbor a diverse community of hydrogenogenic fermenters and hydrogenotrophic methanogens, acetogens, sulfate reducers, fumarate reducers, and denitrifiers. Secondly, we used the model system of the H_2_-producing carbohydrate fermenter *Ruminococcus albus* 7 and the H_2_-utilizing fumarate-reducing syntrophic partner *Wolinella succinogenes* DSM 1740 ^25,53,54,67^ to gain a deeper mechanistic understanding of how and why ruminant bacteria regulate H_2_ metabolism. We observed significant differences in the growth, transcriptome, and metabolite profiles of these bacteria in co-culture compared to pure culture. Finally, we compared gene expression profiles associated with H_2_ metabolism between low-versus high-methane yield sheep ^63^. It was recently proposed, on the basis of community structure analysis, that fewer H_2_-producing bacteria inhabit low methane yield sheep ^68^. In this work, we present an alternative explanation: H_2_ uptake through non-methanogenic pathways accounts for these differences. Whereas the enzymes mediating fermentative H_2_ production are expressed at similar levels, those supporting H_2_ uptake through acetogenesis, fumarate reduction, and denitrification pathways are highly upregulated in low methane yield sheep. In turn, these findings support that strategies to promote alternative H_2_ uptake pathways, including through dietary modulation, may significantly reduce methane emissions.

## Results

### H_2_ metabolism is a common and diverse trait among rumen bacteria, archaea, and eukaryotes

We searched the 501 reference genome sequences of rumen bacteria and archaea ^21^ for genes encoding the catalytic subunits of H_2_-consuming and H_2_-producing enzymes **(Table S1 & S2)**. Of these, 65% encoded the capacity to metabolise H_2_ via [FeFe]-hydrogenases (42%), [NiFe]-hydrogenases (31%), [Fe]-hydrogenases (2.4%), and/or nitrogenases (23%). This suggests that H_2_ metabolism is a widespread trait among rumen microorganisms. We also identified multiple partial sequences of group A1 [FeFe]-hydrogenases in the incomplete genomes of six rumen fungi and ciliates. This is consistent with the known ability of these microorganisms to produce H_2_ during cellulose fermentation ^31^. The 329 hydrogenase- and nitrogenase-positive genomes spanned 108 genera, 26 orders, 18 classes, and 11 phyla **(Figure 1; Figure S1; Table S1 & S2)**.

**Figure 1.**
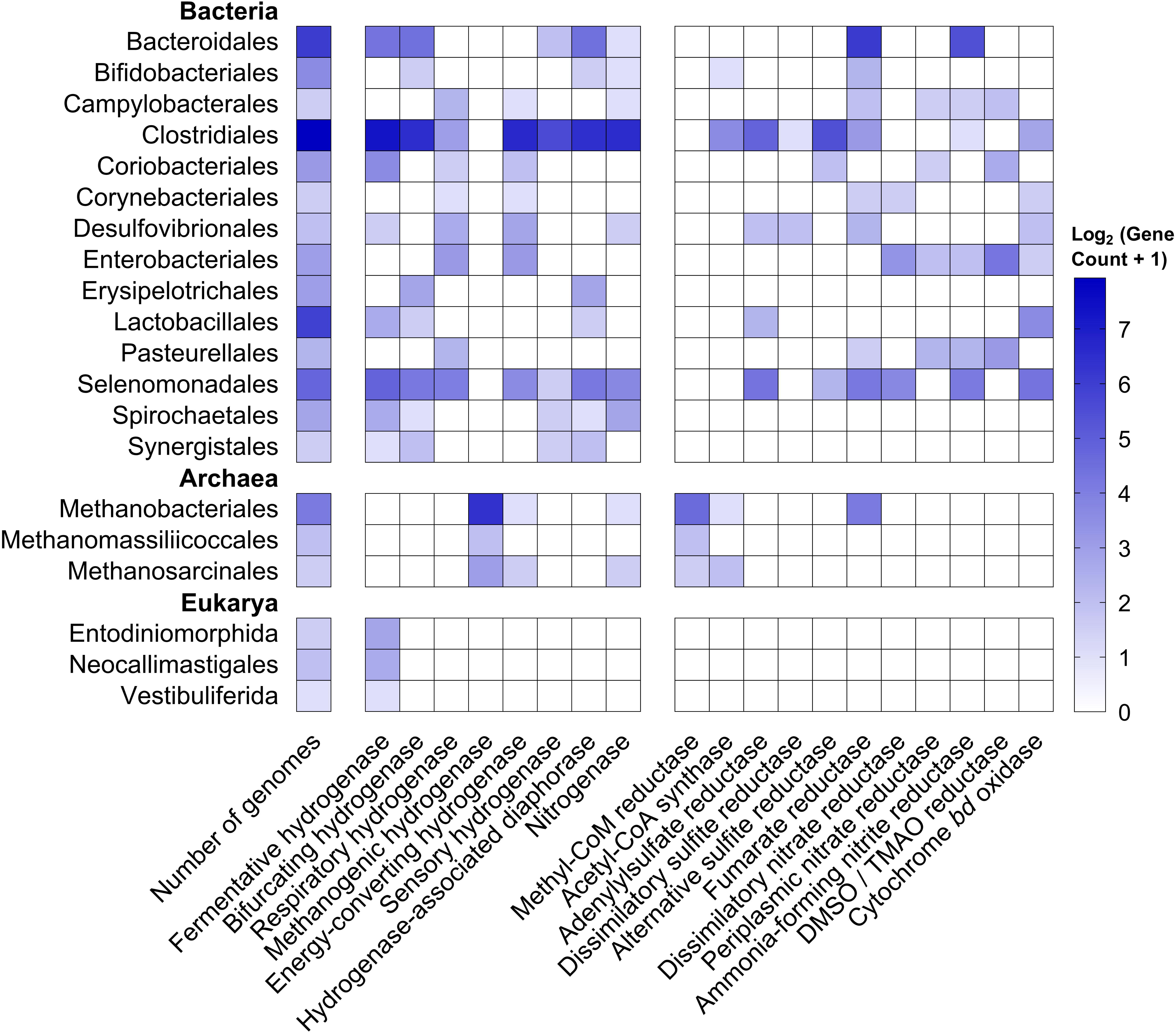
Heatmap showing distribution of enzymes mediating H_2_ production and H_2_ consumption in orders of rumen microorganisms. Results are shown based on screens of the 501 genomes of cultured rumen bacteria and archaea (410 from the Hungate collection plus 91 other genomes). Partial hydrogenase sequences were also retrieved and classified from four rumen ciliates and two rumen fungi. The left-hand side of the heatmap shows the distribution of the catalytic subunits of enzymes that catalyze H_2_ oxidation and production. These are divided into fermentative hydrogenases (H_2_-producing; group A1, A2, B FeFe-hydrogenases), bifurcating hydrogenases (bidirectional; group A3, A4 FeFe-hydrogenases), respiratory hydrogenases (H_2_-uptake; group 1b, 1c, 1d, 1f, 1i, 2d NiFe-hydrogenases), methanogenic hydrogenases (H_2_-uptake; group 1k, 3a, 3c, 4h, 4i NiFe-hydrogenases, Fe-hydrogenases), energy-converting hydrogenases (bidirectional; group 4a, 4c, 4e, 4f, 4g NiFe-hydrogenases), sensory hydrogenases (group C FeFe-hydrogenases), and nitrogenases (H_2_-producing; NifH). The right-hand side shows the distribution of the catalytic subunits of key reductases in H_2_ consumption pathways. They are genes for methanogenesis (McrA, methyl-CoM reductase), acetogenesis (AcsB, acetyl-CoA synthase), sulfate reduction (DsrA, dissimilatory sulfite reductase; AprA, adenylylsulfate reductase; AsrA, alternative sulfite reductase), fumarate reduction (FrdA, fumarate reductase), nitrate ammonification (NarG, dissimilatory nitrate reductase; NapA, periplasmic nitrate reductase; NrfA, ammonia-forming nitrite reductase), dimethyl sulfoxide and trimethylamine *N*-oxide reduction (DmsA, DMSO and TMAO reductase), and aerobic respiration (CydA, cytochrome *bd* oxidase). Only hydrogenase-encoding orders are shown. **Table S2** shows the distribution of these enzymes by genome, **Figure S1** depicts hydrogenase subgroup distribution by class, and **Table S1** lists the FASTA sequences of the retrieved reads.

We then classified the hydrogenases identified into subgroups. To do so, we used the phylogeny-based, functionally-predictive classification scheme of HydDB ^66^, which has been used to understand H_2_ metabolism in a range of organisms and ecosystems ^69–72^. In total, 273 strains encoded hydrogenases from classes that primarily evolve H_2_ under physiological conditions **(Table S2)**. These include group A1 and B [FeFe]-hydrogenases and group 4e [NiFe]-hydrogenases that couple ferredoxin oxidation to H_2_ production in anaerobic bacteria ^73–75^. However, the most widespread hydrogenases are the group A3 [FeFe]-hydrogenases, which were encoded in 43 genera, among them well-characterized carbohydrate fermenters such as *Ruminococcus, Lachnoclostridium*, and *Bacteroides*. These hydrogenases form heterotrimeric complexes, together with diaphorase subunits, that mediate the recently-discovered process of electron-confurcation: coupling co-oxidation of NADH and ferredoxin produced during fermentative carbon degradation to production of H_2_ ^29,76^. This reversible complex can also support hydrogenotrophic acetogenesis ^77^. By retrieving the genes immediately upstream and downstream, we verified that the diaphorase subunits (HydB) of this complex were co-encoded with the retrieved hydrogenase subunits **(Figure 1; Table S2)**.

In addition, multiple organisms encoded hydrogenases and terminal reductases known to support hydrogenotrophic growth **(Figure 1)**. All 21 methanogen genomes surveyed harbored [NiFe]-hydrogenases together with the signature gene of methanogenesis (*mcrA*) **(Figure 1; Table S2)**. These include 14 *Methanobrevibacter* strains, which encoded a complete set of enzymes for mediating hydrogenotrophic methanogenesis through the Wolfe cycle ^8^, including the [Fe]-hydrogenase and the groups 3a, 3c, 4h, and 4i [NiFe]-hydrogenases. Seven genomes encoded both [FeFe]-hydrogenases (A2, A3) and the marker gene for acetogenesis (*acsB*) **(Table S2)**, including known hydrogenotrophic acetogens *Blautia schinkii* ^40^ and *Acetitomaculum ruminis* ^41^. Several subgroups of the group 1 [NiFe]-hydrogenases, all membrane-bound enzymes known to support hydrogenotrophic respiration ^65,78^, were also detected. Most notably, various *Selenomonas, Mitsuokella*, and *Wolinella* strains encoded such hydrogenases together with the signature genes for fumarate reduction (*frdA*) and nitrate ammonification (*narG, napA, nrfA*). As anticipated, the group 1b [NiFe]-hydrogenase and *dsrA* gene characteristic of hydrogenotrophic sulfate reduction were also encoded in the three genomes of ruminal *Desulfovibrio* isolates **(Figure 1; Table S2)**.

### H_2_ is mainly produced by clostridial electron-bifurcating [FeFe]-hydrogenases and consumed by [NiFe]-hydrogenases of methanogens and selenomonads

We then investigated the relative abundance and expression levels of the retrieved hydrogenases in rumen communities. To do so, we used 20 pairs of metagenomes and metatranscriptomes that were previously sequenced from the rumen contents of age- and diet-matched sheep ^63^ **(Table S3)**. Screening these datasets with hydrogenases retrieved from the rumen microbial reference genomes yielded 15,464 metagenome hits (0.015% of all reads) and 40,485 metatranscriptome hits (0.040%) **(Table S4)**. Across the metagenomes, the dominant hydrogenase reads originated from eleven subgroups (A1, A2, A3, B, 3a, 3c, 4e, 4g, 4h, 4i, Fe) **(Figure 2a & S2a)** and three taxonomic orders (Clostridiales, Methanobacteriales, Selenomonadales) **(Figure 2c & S3a)**; this is concordant with the hydrogenase content in the genomes of the dominant community members ^63,64^ **(Table S2)**. Metatranscriptome analysis indicated these genes were differentially expressed: whereas A3, 1d, 3a, 3c, and 4g genes were highly expressed (RNA / DNA expression ratio > 4), others were expressed at moderate (A1, A2, Fe; ratio 1.5 – 2.5) or low levels (B, 4e, 4h, 4i; ratio < 1.5) **(Figure S2 & S3)**. Though putative nitrogenase genes (*nifH*) were detected, expression ratios were low (av. 0.45), suggesting nitrogen fixation is not a significant H_2_ source in sheep **(Figure S4)**.

**Figure 2.**
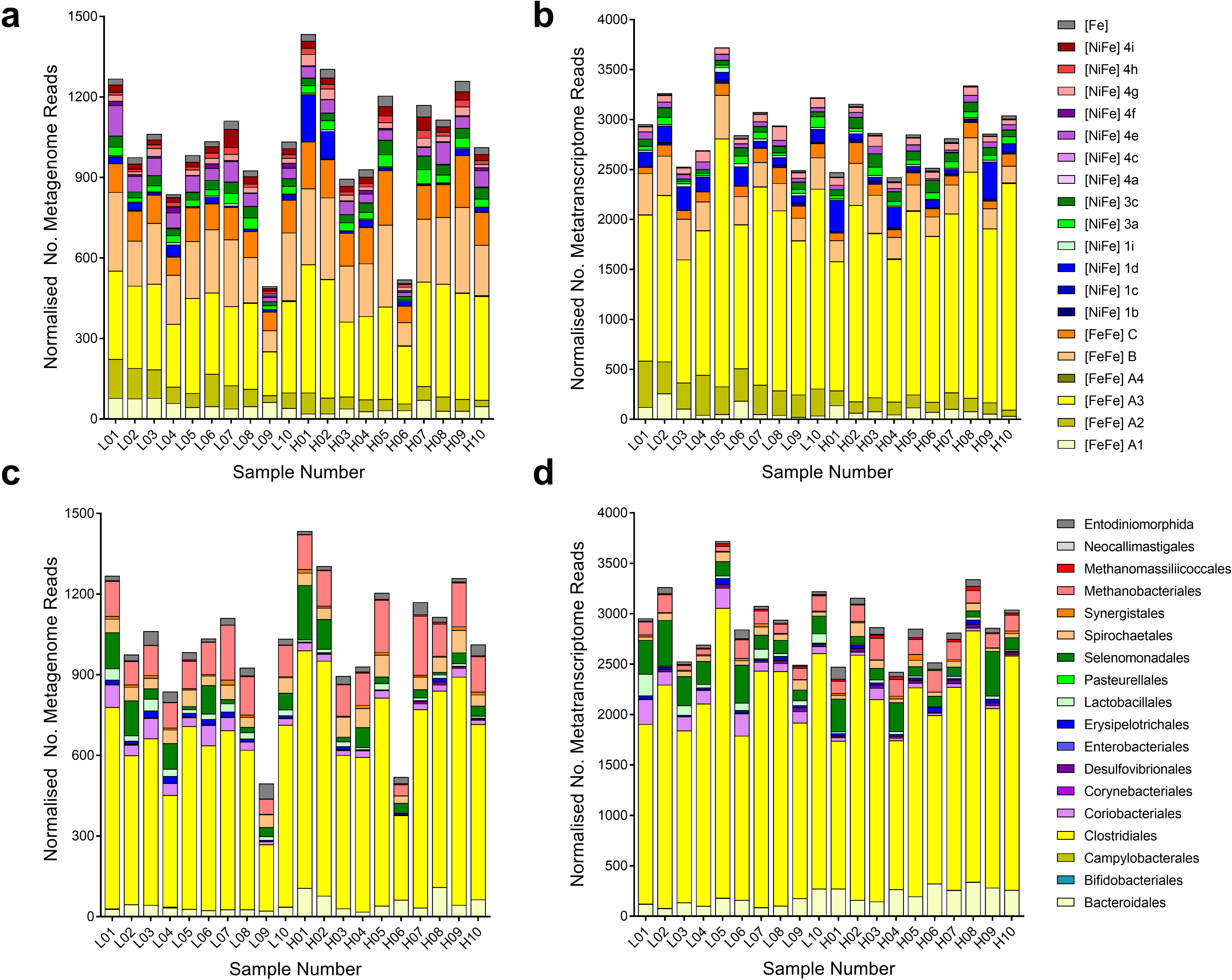
Hydrogenase content in the metagenomes and metatranscriptomes of the microbial communities within rumen contents of high and low methane yield sheep. Hydrogenase content is shown based on hydrogenase subgroup (a, b) and predicted taxonomic affiliation (c, d) for metagenome datasets (a, c) and metatranscriptome datasets (b, d). Hydrogenase-encoding sequences were retrieved from 20 paired shotgun metagenomes and metatranscriptomes randomly subsampled at five million reads. Reads were classified into hydrogenase subgroups and taxonomically assigned at the order level based on their closest match to the hydrogenases within the genomes screened **(Figure 1)**. L01 to L10 are datasets for sheep that were low methane yield at time of sampling, H01 to H20 are datasets from sheep that were high methane yield at time of sampling (see **Table S3** for full details).

Accounting for 54% of hydrogenase transcripts detected **(Figure 2b, 3a, S2)**, group A3 [FeFe]-hydrogenases appear to be the primary catalysts of H_2_ production in ruminants. We assigned the retrieved transcripts to taxa based on their closest hits to the rumen genome hydrogenase dataset **(Table S4)**. Clostridia accounted for the majority of the hits **(Figure 2d)**, including *Ruminococcus* (22%), *Saccharofermentans* (9.2%), and *Lachnoclostridium* (7.4%) species known to fermentatively produce H_2_ ^29,33,79^ **(Figure S5 & S6)**. Transcripts from the characterized fermentative genera *Bacteroides, Butyrivibrio, Clostridium*, and *Sarcina* were also moderately abundant. A further 21% of group A3 [FeFe]-hydrogenase hits were assigned to three uncharacterized cultured lineages within the Clostridia: Clostridiales R-7, Ruminococcaceae P7, and Lachnospiraceae YSB2008 **(Figure S5 & S6)**. This is compatible with our previous studies showing unclassified microorganisms, especially from R-7 group, are abundant in rumen ^21^. H_2_-evolving hydrogenases from the A1 and B subgroups were also detected, but their RNA/DNA expression ratios were threefold lower than the A3 hydrogenases. Rumen ciliates such as *Epidinium* dominated A1 reads **(Figure 2d & S5)**, but it is likely that their abundance in the datasets is underestimated due to the minimal genome coverage of these organisms to date.

**Figure 3.**
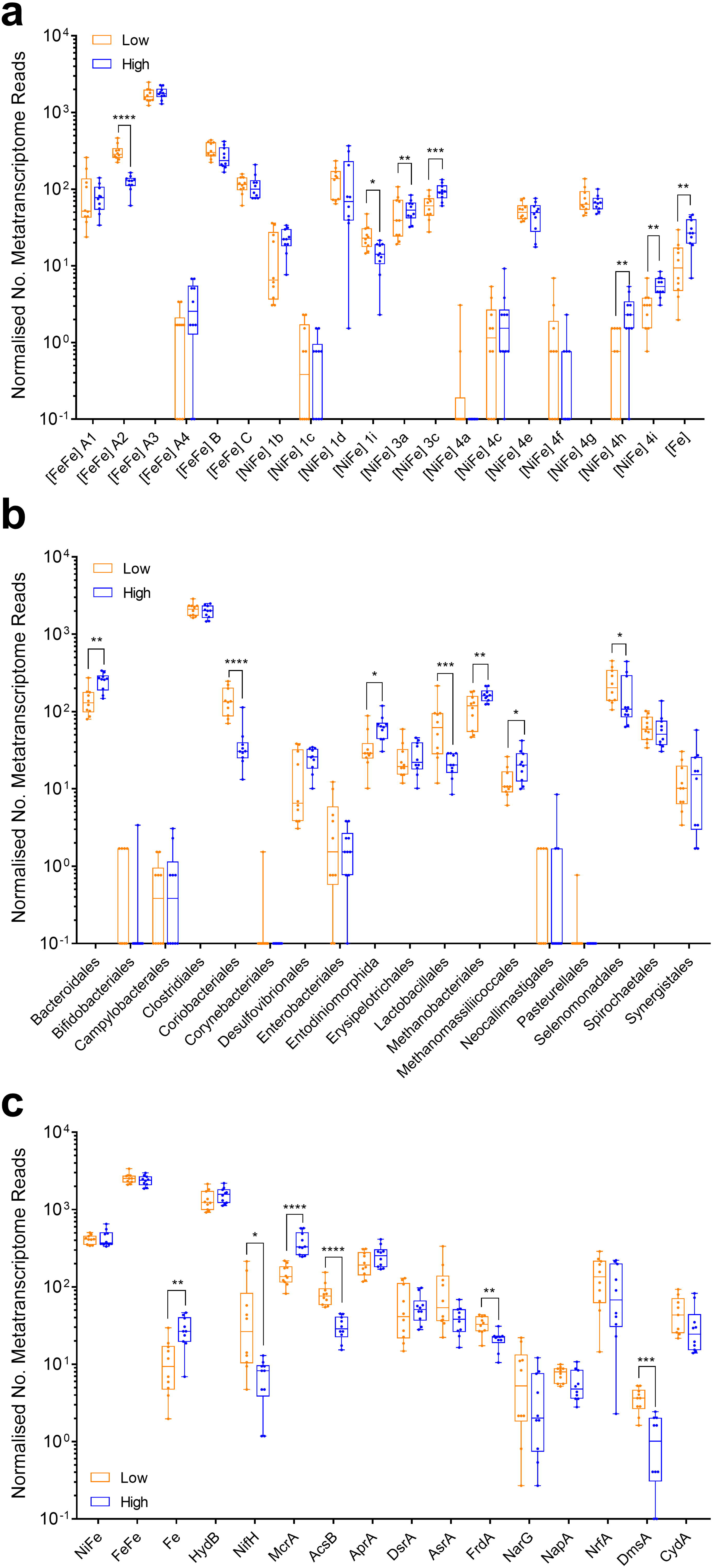
Comparison of expression levels of H_2_ production and H_2_ uptake pathways in low and high methane yield sheep. Results are shown for ten metatranscriptome datasets each from low methane yield sheep (orange) and high methane yield sheep (blue) that were randomly subsampled at five million reads. (a) Normalized count of hydrogenase transcript reads based on hydrogenase subgroup. (b) Normalized count of hydrogenase transcript reads based on predicted taxonomic affiliation. (c) Normalized count of transcript reads of key enzymes involved in H_2_ production and H_2_ consumption, namely the catalytic subunits of [NiFe]-hydrogenases (NiFe), [FeFe]-hydrogenases (FeFe), [Fe]-hydrogenases (Fe), hydrogenase-associated diaphorases (HydB), nitrogenases (NifH), methyl-CoM reductases (McrA), acetyl-CoA synthases (AcsB), adenylylsulfate reductases (AprA), dissimilatory sulfite reductases (DsrA), alternative sulfite reductases (AsrA), fumarate reductases (FrdA), dissimilatory nitrate reductases (NarG), periplasmic nitrate reductases (NapA), ammonia-forming nitrite reductases (NrfA), DMSO / TMAO reductases (DmsA), and cytochrome *bd* oxidases (CydA) are provided. For FrdA, NrfA, and CydA, the numerous reads from non-hydrogenotrophic organisms (e.g. Bacteroidetes) were excluded. Each boxplot shows the ten datapoints and their range, mean, and quartiles. Significance was tested using independent two-group Wilcoxon rank-sum tests (* *p* < 0.05; ** *p* < 0.01; *** *p* < 0.001; **** *p* < 0.0001; full *p* values in **Table S8, Table S9** and **Table S10**). Note the metagenome abundance and RNA / DNA ratio of these genes is shown in **Figure S2** (hydrogenase subgroup), **Figure S3** (hydrogenase taxonomic affiliation), and **Figure S4** (H_2_ uptake pathways). A full list of metagenome and metatranscriptome hits is provided for hydrogenases in **Table S4** and H_2_ uptake pathways in **Table S5.**

The metatranscriptome datasets indicate that multiple H_2_ uptake pathways operate in ruminants **(Figure 2 & 3)**. In agreement with historical paradigms ^6^, hydrogenotrophic methanogenesis appears to be the largest sink of H_2_; methanogens accounted for 5.3% of normalized hydrogenase reads **(Figure 2d)** and methyl-CoM reductase (*mcrA*) is the most expressed of the reductases surveyed **(Figure 3c)**. Consistent with their central roles in the CO_2_-reducing pathway of methanogenesis ^9^, the F_420_-reducing [NiFe]-hydrogenase (3a) ^80^ and the heterodisulfide reductase-associated [NiFe]-hydrogenase (3c) ^81^ of *Methanobrevibacter* species were among the most transcribed of all H_2_ uptake enzymes **(Figure 3a & S5)**. In contrast, the Eha-type (4h), Ehb-type (4i), and [Fe]-hydrogenases were expressed at lower levels **(Figure 3a & S5)**, reflecting their secondary roles in the physiology of methanogens ^82–84^. There was also strong evidence that hydrogenotrophic acetogenesis may be a more significant ruminal H_2_ sink than previously recognized. Across the dataset, acetyl-CoA synthases (*acsB*; 1135 normalized reads) were expressed at a quarter of the level of methyl-CoM reductases (*mcrA*; 5246 normalized reads) **(Figure 3c)**. For 74% of the reads, the closest matches were to predicted hydrogenotrophic acetogens isolated from rumen, including *Blautia, Acetitomaculum*, and *Oxobacter* **(Figure S8 & Table S5)**. Consistently, group A2 and group A3 [FeFe]-hydrogenases from the same genera were moderately expressed in the metatranscriptomes (3.7%) **(Figure S5)**. The other *acsB* reads likely originate from acetogens that use other electron donors, such as formate.

Surprisingly, however, the most highly expressed H_2_ uptake hydrogenase overall is the group 1d [NiFe]-hydrogenase of Selenomonadales (4.1%) **(Figure 3a, 3b & S5)**. This enzyme is likely to mediate the long-known capacity of *Selenomonas* species to grow by hydrogenotrophic fumarate reduction and denitrification ^51,52,57^. Consistently, fumarate reductases (*frdA*), nitrate reductases (*narG*), and ammonia-forming nitrite reductases (*nrfA*) homologous to those in *S. ruminantium* were expressed in the metatranscriptomes **(Figure 3c)**. Normalized *nrfA* expression was fivefold higher than *narG*, suggesting selenomonads may preferentially use external nitrite; while further studies are required to determine the source of nitrite, this compound is known to accumulate in the rumen depending on nitrate content of feed ^85^. Reads corresponding to the group 1b [NiFe]-hydrogenase, periplasmic nitrate reductase (*napA*), *nrfA*, and *frdA* from *Wolinella* was also detected, but at low levels **(Table S4 & S5; Figure S5)**. Several other pathways in low abundance in the metagenome were also highly expressed, notably group 1b [NiFe]-hydrogenases and *dsrA* genes from *Desulfovibrio* species, as well as group 1i [NiFe]-hydrogenases from metabolically flexible Coriobacteriia (e.g. *Slackia, Denitrobacterium*) **(Figure S4 & S5)**. The expression levels of the 1b and 1d hydrogenases, together with the functionally-unresolved 4g hydrogenases, were the highest of all hydrogenases in datasets (RNA / DNA ratio > 10) **(Figure S3)**. Though these findings need to be validated by activity-based studies, they suggest that respiratory hydrogenotrophs are highly active and quantitatively significant H_2_ sinks in the rumen despite often being detected in low abundance ^5^.

### Culture-based studies demonstrate that hydrogenases mediating H_2_ production and uptake are differentially regulated in response to hydrogen levels

In order to better understand how rumen bacteria regulate H_2_ metabolism, we performed a culture-based study using *Ruminococcus albus* 7 and *Wolinella succinogenes* DSM 1740, a model system for interspecies hydrogen transfer ^25^. We compared the growth, transcriptome, and extracellular metabolite profiles of these strains in either pure culture or co-culture when grown on modified fumarate-supplemented Balch medium **(Table S6)**. The concentrations of the metabolites consumed and produced by the strains varied between the conditions **(Table S1; Figure S8)** in a manner consistent with the transcriptomic results **(Figure 4)** and historical paradigms ^24,25,29,53^. Pathway reconstruction indicated that *R. albus* fermentatively degraded cellobiose to H_2_, acetate, and ethanol in pure culture (glucose + 3.3 ADP + 3.3 P_i_ → 2.6 H_2_ + 1.3 acetate + 0.7 ethanol + 2 CO_2_ + 3.3 ATP ^29^) and H_2_ and acetate in co-culture (glucose + 4 ADP + 4 P_i_ → 4 H_2_ + 2 acetate + 2 CO_2_ + 4 ATP ^29^) **(Figure 4a, 4b, 4c)**. *W. succinogenes* grew by hydrogenotrophic fumarate respiration under both conditions by using exogenously supplied H_2_ in pure culture and syntrophically-produced H_2_ in co-culture **(Figure 4d, 4e, 4f)**. Hence, *R. albus* channels fermentation through the pathway that yields stoichiometrically more ATP, H_2_, and acetate, provided that H_2_ concentrations are kept sufficiently low through interspecies hydrogen transfer for this to be thermodynamically favorable.

**Figure 4.**
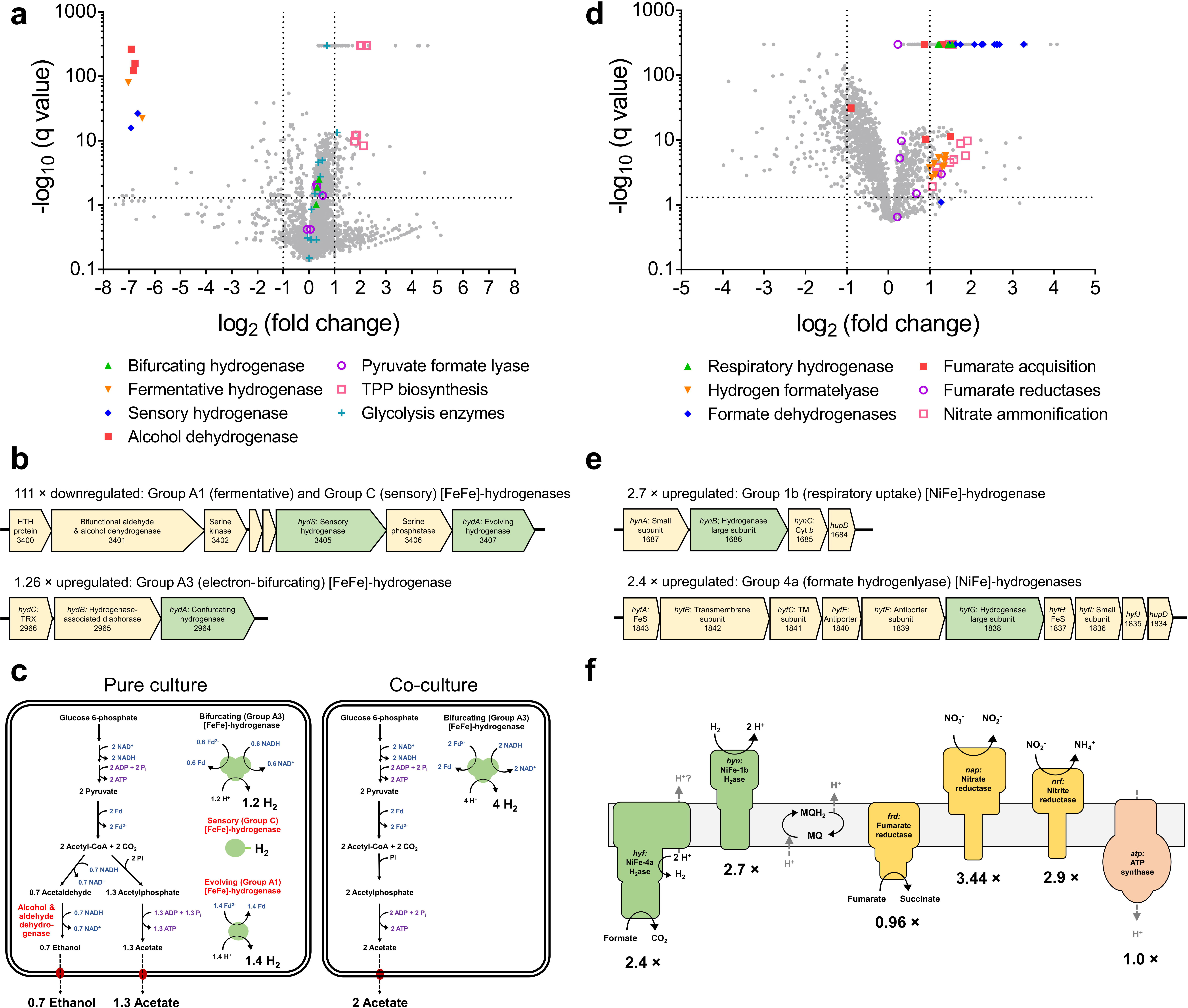
Comparison of whole genome expression levels of *Ruminococcus albus* and *Wolinella succinogenes* in pure culture and co-culture. Pure cultures and co-cultures of *Ruminococcus albus* 7 (a, b, c) and *Wolinella succinogenes* DSM 1740 (d, e, f) were harvested in duplicate during mid-exponential phase and subject to RNA sequencing. (a & d) Volcano plots of the ratio of normalized average transcript abundance for co-cultures over pure cultures. Each gene is represented by a grey dot and key metabolic genes, including hydrogenases, are highlighted as per the legend. (b & d) Predicted operon structure of the three hydrogenases of *R. albus* and two hydrogenases of *W. succinogenes*. (e) Comparison of dominant fermentation pathways of *R. albus* in pure culture (left) and co-culture (right) based on transcriptome reads and metabolite profiling. The three enzymes downregulated in co-culture are in red font. (f) Respiratory chain composition of *W. succinogenes* in pure culture and co-culture based on transcriptome reads. Metabolite profiling indicated that the respiratory hydrogenase and fumarate reductases were active in both conditions. A full list of read counts and expression ratios for each gene is provided in **Table S**

Transcriptome profiling revealed that *R. albus* tightly regulates the expression of its three hydrogenases **(Figure 4a, 4b)**. Overall, 133 genes were differentially expressed (fold change > 2, *q*-value < 0.05) in co-culture compared to pure culture **(Table S7)**. Of these, the greatest fold-change was the 111-fold downregulation of a putative eight-gene cluster encoding the ferredoxin-only hydrogenase (group A1 [FeFe]-hydrogenase), a bifunctional alcohol and aldehyde dehydrogenase, and regulatory elements including a putative sensory hydrogenase (group C [FeFe]-hydrogenase) **(Figure 4a & 4b)**. By suppressing expression of these enzymes, *R. albus* can divert carbon flux from ethanol production to the more energetically efficient pathway of acetate production; acetate fermentation produces equimolar levels of NADH and reduced ferredoxin, which can be simultaneously reoxidized by the electron-bifurcating hydrogenase (group A3 [FeFe]-hydrogenase) **(Figure 4c)**. Glycolysis enzymes and the phosphate acetyltransferase, acetate kinase, and electron-bifurcating hydrogenase of the acetate production pathway were expressed at similarly high levels under both conditions **(Figure 4a & 4b)**. However, there was a significant increase in the biosynthesis of thiamine pyrophosphate, a cofactor for pyruvate dehydrogenase complex, in co-culture **(Figure 4a)**.

The fermentation stoichiometries of *R. albus* 7 measured in pure culture compared to co-culture **(Table 1)** were the same as we previously reported for the bacterium at high vs low concentrations of H_2_ ^29^. This suggests that the differences in regulation are primarily determined by H_2_ levels, rather than by direct interactions with syntrophic partners. This regulation may be achieved through direct sensing of H_2_ by the putative sensory group C [FeFe]-hydrogenase co-transcribed with the ferredoxin-only hydrogenase and alcohol dehydrogenase **(Figure 4e)**. In common with other enzymes of this class ^65,86,87^, this enzyme contains a H-cluster for H_2_ binding, a PAS domain for signal transfer, and a putative serine or threonine phosphatase that may modify downstream regulators. Thus, analogous to the well-studied regulatory hydrogenases of aerobic bacteria ^88,89^, this enzyme may directly sense H_2_ levels and induce expression of the alcohol / aldehyde dehydrogenase and ferredoxin-only hydrogenase when H_2_ concentrations are high through a feedback loop. H_2_ sensing may be a general mechanism regulating hydrogenase expression in ruminants, given group C [FeFe]-hydrogenases are abundant in ruminant genome **(Figure 1)**, metagenome **(Figure 2a)**, and metatranscriptome datasets **(Figure 2b & (3a)**.

**Table 1.** Comparison of growth parameters and metabolite profiles of *Ruminococcus albus* and *Wolinella succinogenes* in pure culture and co-culture. Growth of pure cultures and co-cultures of *Ruminococcus albus* 7 and *Wolinella succinogenes* DSM 1740 was monitored by qPCR. Values show means ± standard deviations of three biological replicates. Also shown is the change in extracellular pH, percentage hydrogen gas (measured by gas chromatography), and concentrations of fumarate, succinate, acetate, ethanol, and formate (measured by ultra-fast liquid chromatography) between 0 hours and 12 hours. Growth media was the same between the three conditions, except 80% H_2_ was added for *W. succinogenes* growth, whereas no H_2_ was added for the other conditions. Full liquid metabolite measurements are shown in **Figure S8**. BDL = below detection limit.

|  | *Ruminococcus albus* | *Wolinella succinogenes* | Co-culture |
| --- | --- | --- | --- |
| **Growth parameters** |  |  |  |
| Growth yield (OD<sub>600</sub>) | 0.79±0.01 | 0.36±0.01 | 0.93±0.01 |
| Specific growth rate (h<sup>-1</sup>) | 0.58±0.19 | 0.33±0.06 | 0.57±0.34 (*Ra*)<br>0.54±0.11 (*Ws*) |
| **Concentration changes of extracellular metabolites** |  |  |  |
| Hydrogen (%) | +5.3 | -78.4 | BDL |
| Fumarate (mM) | -5.5 | -46.3 | -43.1 |
| Succinate (mM) | +2.2 | +54.6 | +55.4 |
| Acetate (mM) | +21.8 | 0 | +32.4 |
| Ethanol (mM) | +8.7 | 0 | +0.3 |
| Formate (mM) | BDL | BDL | BDL |
| pH | -0.4 | -0.4 | -0.6 |

The transcriptome results also clarified understanding of hydrogenotrophic fumarate respiration by *W. succinogenes* **(Figure 4d)**. In both pure culture and co-culture, the group 1b [NiFe]-hydrogenase, fumarate reductase, and F_1_F_o_-ATPase that mediate this process were expressed at high levels **(Table S7; Figure 4f)**. A periplasmic asparaginase, aspartate ammonia-lyase, and dicarboxylate-binding proteins were also highly expressed; this suggests that the organism can efficiently produce and import additional fumarate from amino acid sources **(Table S7)**. In total, 352 genes were significantly differentially regulated in co-culture (fold-change > 2, q-value < 0.05). The respiratory hydrogenase was among the upregulated genes **(Figure 4e)**, which may reflect the strain’s faster growth rate in co-culture **(Table 1)**. The periplasmic nitrate reductase and ammonia-forming nitrite reductase **(Figure 4f)** were also induced, indicating some plasticity in oxidant usage, in line with the metatranscriptomic findings. Two formate dehydrogenases and a formate hydrogenlyase (group 4a [NiFe]-hydrogenase) were highly expressed in co-culture **(Figure 4d & 4e)**. This suggests the bacterium can potentially use formate, known to be produced through formate pyruvate lyase by *R. albus*, as an additional electron donor **(Figure 4a)**. However, the significance of these findings is unclear given no formate was detected under any condition **(Table 1)** and the expression of formate-dependent hydrogenases was extremely low in metatranscriptome datasets **(Figure 3c)**.

### Hydrogenotrophic acetogenesis, fumarate reduction, and denitrification pathways are significantly upregulated in low methane yield ruminants

Finally, we tested whether the abundance and expression of hydrogenases and H_2_ uptake pathways differed between high and low methane yield sheep. The current leading hypothesis, proposed on the basis of community composition ^68^, asserts that H_2_ production levels account for differences in methane yield between sheep. To the contrary, the expression levels of the dominant H_2_-evolving hydrogenases (e.g. group A3 [FeFe]-hydrogenases) and taxonomic orders (e.g. Clostridiales) were in fact extremely similar between the groups **(Figure 3a & 3b; Table S8 & S9)**.

We therefore formulated an alternative hypothesis: H_2_ utilization through non-methanogenic pathways can reduce methane yield. In line with this hypothesis, the expression levels of the five methanogen hydrogenases and methyl-CoM reductase are significantly reduced in low methane yield sheep **(Figure 3a & 3c; Table S8 & 10)**, confirming a strong correlation with measured phenotypes **(Table S3)**. Concurrent increases in the gene expression for two major alternative H_2_ sinks were detected, namely acetogenesis (*acsB*; *p* < 0.0001) and fumarate reduction (*frdA*; *p* = 0.002) **(Figure 3c; Table S10)**, concomitant with significant increases in the expression levels of *Blautia* and *Selenomonas* hydrogenases **(Figure S5)**. Expression levels of *nrfA* were also on average 1.8-fold higher in low methane yield sheep, though there was much inter-sample variation in the read count for this gene. Whereas there are more transcripts of *mcrA* than other terminal reductases combined in high methane yield sheep, the transcript levels of *acsB* and *nrfA* together exceed those of *mcrA* in low methane yield sheep. Depending to what extent expression levels predict activity, hydrogenotrophic acetogens and selenomonads may therefore be more active than methanogens in low methane yield sheep and may significantly limit substrate supply for methanogenesis. Two other potential H_2_ sinks are also upregulated in the low methane yield sheep: the putative group 1i [NiFe]-hydrogenase of Coriobacteriia and, consistent with previous observations ^64^, the functionally-unresolved group A2 [FeFe]-hydrogenase of *Sharpea, Olsenella*, and *Oribacterium* **(Figure 3a, 3b, S5)**.

## Discussion

To summarize, H_2_ metabolism is a more widespread and complex process in ruminants than previously realized. Together, the genomic, metagenomic, and metatranscriptomic surveys suggest that multiple orders of bacteria, archaea, and eukaryotes encode and express enzymes mediating H_2_ production and consumption in the rumen. We infer that fermentative Clostridia are the main source of H_2_ in the rumen, which largely agrees with findings from activity-based and culture-based studies ^6,25,26,29^. However, a surprising finding is that uncharacterized lineages within the Clostridia account for a large proportion of hydrogenase reads, emphasizing the need for physiological and bacteriological characterization of these organisms. Further studies are also needed to better account for the role of rumen ciliates and fungi, which to date are underrepresented in genomic datasets.

One of the most important findings of this work is that the recently-characterized electron-bifurcating hydrogenases appear to primarily mediate ruminal H_2_ production. These enzymes are highly upregulated compared to ferredoxin-only hydrogenases *in situ* and constitute over half of hydrogenase reads in metatranscriptomes. We provide a rationale for this finding by showing that *Ruminococcus albus*, a dominant H_2_ producer within the rumen, expresses its electron-bifurcating hydrogenase and suppresses its ferredoxin-only hydrogenase when grown syntrophically with *Wolinella succinogenes*. In this condition, H_2_ concentrations remain sufficiently low that the fermentation pathway producing higher levels of ATP, H_2_, and acetate remains thermodynamically favorable. In the rumen, where tight coupling of hydrogenogenic and hydrogenotrophic processes usually keeps H_2_ at sub-micromolar concentrations ^45^, Clostridia will also preferentially oxidize carbohydrates through higher ATP-yielding pathways and reoxidize the NAD and ferredoxin reduced using the electron-bifurcating hydrogenase. It is likely that the ferredoxin-only hydrogenases are preferentially upregulated during the transient periods where H_2_ levels are high, for example immediately after feeding ^45^. Based on these findings and previously published results ^29,65,86,87^, we propose that the hydrogenases and fermentation pathways are differentially regulated as a result of direct H_2_ sensing by putative sensory [FeFe]-hydrogenases.

The other major finding of this work is that there are multiple highly active H_2_ sinks in the rumen. We provide evidence, based on transcript levels of their hydrogenases and terminal reductases, that acetogens (*Blautia, Acetitomaculum*), fumarate reducers and denitrifiers (*Selenomonas, Wolinella*), and sulfate reducers (*Desulfovibrio*) are quantitatively significant H_2_ sinks in sheep. In support of these findings, our culture-based study confirmed that the enzymes mediating hydrogenotrophic fumarate reduction and potentially nitrate ammonification are highly expressed by *W. succinogenes* in co-culture with *R. albus*. While alternative H_2_ uptake pathways have been previously detected *in vitro* ^40,41,49–52,56,57^, it has generally been assumed that they are quantitatively insignificant compared to hydrogenotrophic methanogenesis ^5,6,45^. To the contrary, hydrogenase and terminal reductase transcripts from alternative H_2_ uptake pathways are more numerous than those of methanogens in low methane yield sheep, and hence these pathways may collectively serve as a larger H_2_ sink than methanogenesis under some circumstances. These findings justify activity-based studies to quantity H_2_ flux within ruminants between the pathways. There is also evidence of other novel pathways operating in the rumen, mediated by the functionally unresolved group 1i [NiFe]-hydrogenases (*Slackia, Denitrobacterium*), group 4g [NiFe]-hydrogenase (*Clostridium*), and group A2 [FeFe]-hydrogenases (*Sharpea, Oribacterium, Olsenella*).

The strong correlation between H_2_ uptake pathways and methane yield phenotypes suggests that modulating H_2_ metabolism may be an effective methane mitigation strategy. One strategy is to develop inhibitors that redirect electron flux from H_2_ production towards volatile fatty acid production. However, given the central role of H_2_ metabolism in the physiology and ecology of most rumen microorganisms, this would be challenging to achieve without compromising rumen function and consequently ruminant nutrition. Furthermore, such strategies may have a converse effect on methane production, given lower H_2_ concentrations restrict acetogens more than methanogens ^45^. Instead, our metatranscriptome analyses suggest a more promising approach may be to stimulate alternative H_2_ pathways such as fumarate, nitrate, and sulfate respiration. Selective breeding of low methane yield sheep is an option, given methane yield and in turn metatranscriptome profiles have been shown to be a quantitative hereditable trait to some extent ^63,90^. However, the similar metagenome profiles between the sheep, combined with the metatranscriptome profiles of the phenotype-switching sheep, indicate alternative H_2_ uptake pathways are also inducible. Another solution may be to supplement animal feeds with electron acceptors, such as fumarate, nitrate, or sulfate, that stimulate the dominant respiratory hydrogenotrophs. Such approaches have shown some promise in mitigating methane production both *in vitro* ^91–93^ and in field trials ^61,62,94,95^. These strategies may complement methanogenesis inhibitors ^16,17^ by facilitating the redirection of H_2_ flux from methanogens to other pathways.

## Materials and Methods

### Comparative genomic analysis

The protein sequences of the 501 genomes of cultured rumen bacteria (410 from Hungate Collection ^21^, 91 from other sources) were retrieved from the Joint Genome Institute (JGI) genome portal. These sequences were then screened against local protein databases for the catalytic subunits of the three classes of hydrogenases (NiFe-hydrogenases, FeFe-hydrogenases, Fe-hydrogenases), nitrogenases (NifH), methyl-CoM reductases (McrA), acetyl-CoA synthases (AcsB), adenylylsulfate reductases (AprA), dissimilatory sulfite reductases (DsrA), alternative sulfite reductases (AsrA), fumarate reductases (FrdA), dissimilatory nitrate reductases (NarG), periplasmic nitrate reductases (NapA), ammonia-forming nitrite reductases (NrfA), DMSO / TMAO reductases (DmsA), and cytochrome *bd* oxidases (CydA). Hydrogenases were screened using the HydDB dataset ^66,96^, targeted searches were used to screen six protein families (AprA, AsrA, NarG, NapA, NrfA, DmsA, CydA), and comprehensive custom databases were constructed to screen five other protein families (NifH, McrA, AcsB, DsrA, FrdA) based on their total reported genetic diversity ^97–101^. A custom Python script incorporating the Biopython package ^102^ for producing and parsing BLAST results was used to batch-submit the protein sequences of the 501 downloaded genomes as queries for BLAST searches against the local databases. Specifically, hits were initially called for alignments with an e-value threshold of 1e-50 and the resultant XML files were parsed. Alignments producing hits were further filtered for those with coverage values exceeding 90% and percent identity values of 30% to 70%, depending on the target, and hits were subsequently manually curated. **Table S1** provides the FASTA protein sequences and alignment details of the filtered hits. For hydrogenases, the protein sequences flanking the hydrogenase large subunits were also retrieved; these sequences were used to classify group A [FeFe]-hydrogenases into subtypes (A1 to A4), as previously described ^96^, and retrieve diaphorase sequences (HydB) associated with the A3 subtype. Partial [FeFe]-hydrogenase protein sequences from six incompletely sequenced rumen ciliates and fungi genomes were retrieved through targeted blastP searches ^103^ in NCBI.

### Metagenomic and metatranscriptomic analysis

We analyzed previously published datasets of twenty paired metagenomes and metatranscriptomes of sheep rumen contents ^63^. All profiles were derived from the rumen contents of age-matched, pelleted lucerne-fed rams that were collected four hours after morning feeding and subject to paired-end sequencing on the HiSeq 2000 platform ^63^. The samples were taken from ten rams at two different sampling dates based on their measured methane yields ^63,90^; four rams were consistently low yield, four were consistently high yield, and two others switched in methane yield between the sampling dates **(Table S3)**. The metagenome and metatranscriptome datasets analyzed are accessible at the NCBI Sequence Read Archive (SRA; http://www.ncbi.nlm.nih.gov/sra) accession numbers SRA075938, and SRX1079958 - SRX1079985 under bioproject number PRJNA202380. Each metagenome and metatranscriptome was subsampled to an equal depth of 5 million reads using seqtk (https://github.com/lh3/seqtk) seeded with parameter -s100. Subsampled datasets were then screened in DIAMOND (default settings, one maximum target sequence per query) ^104^ using the protein sequences retrieved from the 507 rumen microbial genomes (NiFe-hydrogenases, FeFe-hydrogenases, Fe-hydrogenases, HydB, NifH, McrA, AcsB, AprA, DsrA, AsrA, FrdA, NarG, NapA, NrfA, DmsA, CydA). Results were then filtered (length of amino acid > 40 residues, sequence identity > 65%). Subgroup classification and taxonomic assignment of the hydrogenase reads was based on their closest match to the hydrogenase dataset derived from the 507 genomes at either 65% or 85% identity. The number of reads with the rumen-specific hydrogenase dataset (15464 metagenome hits, 40485 metatranscriptome hits) exceeded those obtained by screening with the generic dataset from HydDB ^66^ (12599 metagenome reads, 31155 metatranscriptome reads), verifying the rumen dataset comprehensively captures hydrogenase diversity. For each dataset, read count was normalized to account for the average length of each gene using the following formula: Normalized Read Count = Actual Read Count × (1000 / Average Gene Length). Independent two-group Wilcoxon rank-sum tests were used to determine whether there were significant differences in the targets analyzed between low and high methane yield sheep. Separate analyses were performed based on gene abundance, transcript abundance, and RNA/DNA ratio.

### Bacterial growth conditions and quantification

The bovine rumen isolates *Ruminococcus albus* 7 ^67^ and *Wolinella succinogenes* DSM 1740 ^53^ were cultured anaerobically at 37°C in modified Balch medium ^105^ **(Table S6)**. Pre-cultures were grown in Balch tubes (18 × 150 mm; Chemglass Life Sciences, Vineland, NJ) containing 20% v/v culture medium and sealed with butyl rubber stoppers crimped with aluminium caps. Cultures were grown in Pyrex side-arm flanks (Corning Inc., Corning, NY) containing 118 mL modified Balch medium. Two pre-cultures were grown before final inoculation, and all inoculum transfers were 5% (v/v). The headspace consisted of 20% CO_2_ and 80% N_2_ for *R. albus* pure cultures and the co-cultures, and 20% CO_2_ and 80% H_2_ for *W. succinogenes* pure cultures. Cultures were periodically sampled at 0, 3, 5, 7, 9, and 11 h for metabolite analysis and bacterial quantification. Culture samples were immediately centrifuged (16,000 × *g*, 10 min) in a bench-top centrifuge (Eppendorf, Hamburg, Germany). For metabolite analysis, the supernatant was collected and further centrifuged (16,000 × *g*, 10 min) before HPLC analysis. For bacterial quantification, DNA was extracted from each pellet using the Fungal/Bacterial DNA MiniPrep kit according to the manufacturer’s instructions (Zymo Research, Irvine, CA). Quantitative PCR (qPCR) was used to quantify the number of copies of the Rumal_2867 (*R. albus* glucokinase gene; FW: CTGGGATTCCTGAACTTTCC; RV: ATGCATACTGCGTTAG) and WS0498 (*W. succinogenes flgL* gene; FW: CAGACTATACCGATGCAACTAC; RV: GAGCGGAGGAGATCTTTAATC) against pGEM-T-Easy standards of each gene of known concentration. DNA was quantified using the iTaq Universal SYBR Green Mix (Bio-Rad) using a LightCycler 480 (Roche Holding AG, Basel, Switzerland).

### Liquid and gas metabolite analysis

The concentrations of acetate, ethanol, fumarate, succinate, and formate in the culture supernatants were analyzed using an Ultra-Fast Liquid Chromatograph (UFLC; Shimadzu, Kyoto, Japan). The UFLC consisted of a DGU-20A5 degasser, a SIL-20ACHT autosampler, an LC-20AT solvent delivery unit, an RID-10A refractive index detector, a CBM-20A system controller, and a CTO-20AC column oven. The mobile phase was 5 mM H_2_SO_4_ passed through an Aminex HPX-87H ion exclusion column (Bio-Rad, Hercules, CA) at a flow rate of 0.4 mL min^-1^, 25°C. Each culture was also sampled at 0 h and 24 h to analyze H_2_ percentage mixing ratios using a gas chromatograph (GC; Gow-Mac Series 580 Thermal Conductivity Gas Chromatograph, Gow-Mac Instrument Co., Bethlehem, PA). Samples were withdrawn directly from the culture tube in a gas-tight syringe and 0.5 mL was injected into GC for analysis using N_2_ as the carrier gas. The flow rate was 60 mL min^-1^, the detector was set to 80°C, the injector was set to 80°C, and the oven was set to 75°C. For both liquid and gas analyses, peak retention times and peak area were compared to standards of known concentration.

### RNA extraction and sequencing

Each pure culture and co-culture used for transcriptome analysis was grown in duplicate in Balch tubes. Growth was monitored until the cultures were in mid-exponential phase; the change in OD_600_ at this phase was 0.14 for *W. succinogenes*, 0.20 for *R. albus*, and 0.35 for the co-culture. At mid-exponential phase, 5 mL cultures were harvested by centrifugation (13,000 x *g*, 4°C). Cell pellets were resuspended in 400 μL fresh lysis buffer (5 mM EDTA, 0.5% SDS, 25 mM lysozyme, 250 U mL^-1^ mutanolysin, and 150 μg mL^-1^ proteinase K in 25 mM sodium phosphate buffer, pH 7.0) and incubated 30 minutes at 55°C with periodic vortexing. RNA was subsequently extracted using an RNeasy Mini Kit following the manufacturer’s protocol, including all optional steps (Qiagen, Hilden, Germany) and eluted with 50 μL ultra-pure DEPC-treated water (Invitrogen, Carlsbad, CA). RNA quantity, quality, and integrity were confirmed by Qubit Fluorometry (Invitrogen, Carlsbad, CA), Nanodrop UV-Vis Spectrophotometry (Thermo Fisher Scientific, model 2300c), and agarose gel electrophoresis respectively. Bacterial rRNA was removed from 1 μg of total RNA with the MicrobExpress Kit (Life Technologies, Carlsbad, CA). Libraries were prepared on the enriched mRNA fraction using the Tru-Seq Stranded RNA Sample Prep Kit (Illumina, San Diego, CA). The barcoded libraries were pooled in equimolar concentration the pool and sequenced on one lane for 101 cycles on a HiSeq2000 using a TruSeq SBS Sequencing Kit (Version 3). Fastq files were generated and demultiplexed with the bc12fastq Conversion Software (Illumina, version 1.8.4). The RNA-seq data were analyzed using CLC Genomics Workbench version 5.5.1 (CLC Bio, Cambridge, MA). RNA-seq reads were mapped onto the reference genome sequences of *Ruminococus albus* 7 ^106^ and *Wolinella succinogenes* DSM 1740 ^107^ **(Table S7 & S11)**. The RNA-seq output files were analyzed for statistical significance as described ^108^ and q-values were generated using the qvalue package in R ^109^. Predicted subsystems and functions were downloaded and aligned to the RNA-seq transcriptional data using the RAST Server ^110^.

## Supporting information

## Acknowledgements

This study was funded by the New Zealand Government to support the objectives of the Livestock Research Group of the Global Research Alliance on Agricultural Greenhouse Gases *via* a grant from the New Zealand Fund for Global Partnerships in Livestock Emissions Research (SOW14-GPLER4-AGR-SP6; awarded to G.T.A., S.C.L., W.J.K., R.M., S.K., and G.M.C.). Transcriptomic and metabolic research on the co-cultures was supported by the Agriculture and Food Research Initiative competitive grant 2012-67015-19451 from the USDA National Institute of Food and Agriculture (awarded to R.I.M. and I.C.). The study was also supported by an ARC DECRA Fellowship (DE170100310; awarded to C.G.), an ARC Future Fellowship (FT170100441; awarded to M.J.M.), and PhD scholarships awarded by the University of Otago (C.W.) and Monash University (L.C.W.).

## Author contributions

G.M.C., R.I.M., G.T.A., I.C., S.C.L., W.J.K., S.K., and C.G. conceived this study. C.G., R.I.M., G.M.C., I.C., S.E.M., M.J.M., and X.C.M. designed research, supervised students, and analyzed data. C.G., L.C.W., M.J.M., and S.C.L. performed the comparative genomic analysis. C.G., C.W., S.E.M., G.M.C., R.R.G., G.T.A., W.J.K., S.C.L., and X.C.M. performed the metagenomic and metatranscriptomic analysis. R.G. performed the co-culture experiments and R.G., R.I.M., I.C., and C.G. analyzed the results. C.G. wrote and illustrated the paper with input from all authors.

The authors declare no conflict-of-interest.

