## Supplementary material for "Alternative hydrogen uptake pathways suppress methane production in ruminants"

**Supplementary Information**

**Table S1 (xlsx).** Protein FASTA sequences of the enzyme subunits retrieved from the genomes of 501 rumen bacteria and archaea. The retrieved sequences of the catalytic subunits of the [NiFe]-hydrogenases (NiFe), [FeFe]-hydrogenases (FeFe), [Fe]-hydrogenases (Fe), hydrogenase-associated diaphorase (HydB), nitrogenases (NifH), methyl-CoM reductases (McrA), acetyl-CoA synthases (AcsB), adenylylsulfate reductases (AsrA), dissimilatory sulfite reductases (DsrA), anaerobic sulfite reductases (AsrA), fumarate reductases (FrdA), dissimilatory nitrate reductases (NarG), periplasmic nitrate reductases (NapA), ammonia-forming nitrite reductases (NrfA), DMSO / TMAO reductases (DmsA), and cytochrome *bd* oxidases (CydA) are provided.

**Table S2 (xlsx).** Taxonomic distribution of H_2_-metabolizing enzymes and H_2_ uptake pathways in 501 genomes of rumen-associated bacteria and archaea.

**Table S3 (xlsx).** Details of the metagenome and metatranscriptome samples obtained from the rumen of low and high methane yield sheep. The sheep ID, sampling date, methane yield, and methane rank for each pair of metagenome and metatranscriptome samples is included.

**Table S4 (xlsx).** List of hydrogenase hits retrieved from the metagenome and metatranscriptome datasets of low and high methane yield sheep rumen. Hits can be sorted and filtered by microbial taxonomy, sheep rank, and hydrogenase subgroup.

**Table S5 (xlsx).** List of non-hydrogenase hits retrieved from the metagenome and metatranscriptome datasets of low and high methane yield sheep rumen. Hits can be sorted and filtered by microbial taxonomy, sheep rank, and enzyme family.

**Table S6 (xlsx).** Composition of the modified Balch medium used for culturing *Ruminococcus albus* and *Wolinella succinogenes* in pure culture and co-culture.

**Table S7 (xlsx).** Transcript read count for RNA-seq experiment comparing gene expression of *Ruminococcus albus* 7 and *Wolinella succinogenes* DSM 1740 in pure culture and co-culture.

**Table S8.** Significance testing of the differences in gene and transcript abundance of hydrogenase subgroups observed in the rumen of low and high methane yield sheep. *p* values are shown for differences in abundance of each gene class in the metagenome (column 3), the metatranscriptome (column 4), and the ratio between them (column 5). In all cases, significance was tested using independent two-group Wilcoxon rank-sum tests.

| **[FeFe]-hydrogenases** | | **Gene** | **Transcript** | **Ratio** |
| --- | --- | --- | --- | --- |
| [FeFe] A1 | Evolving: Ferredoxin-coupled | 0.002 (**) | 0.342 | 0.012 (*) |
| [FeFe] A2 | Putative: Glutamate synthase-linked | 0.0004 (***) | <0.0001 (****) | 0.631 |
| [FeFe] A3 | Bifurcating: NADH and ferredoxin dependent | 0.060 | 0.436 | 0.578 |
| [FeFe] A4 | Bifurcating: Formate dehydrogenase-linked | 0.721 | 0.160 | N/A |
| [FeFe] B | Evolving: Colonic-type | 0.448 | 0.424 | 0.631 |
| [FeFe] C | Sensory | 0.063 | 0.578 | 0.143 |
| HydB | [FeFe] A3 diaphorase subunit | 0.015 (*) | 0.247 | 0.631 |
| **[NiFe]-hydrogenases** | | **Gene** | **Transcript** | **Ratio** |
| [NiFe] 1b | Uptake: Prototypical | 0.069 | 0.109 | 0.0004 (***) |
| [NiFe] 1c | Uptake: Hyb-type | 1 | 0.913 | N/A |
| [NiFe] 1d | Uptake: Oxygen-tolerant | 1 | 0.210 | 0.029 (*) |
| [NiFe] 1i | Uptake: Coriobacteria-type | 0.988 | 0.037 (*) | 0.667 |
| [NiFe] 3a | Uptake: F_420_-reducing | 0.158 | 0.001 (**) | 0.052 |
| [NiFe] 3c | Uptake: Heterodisulfide-reducing | 0.541 | 0.0005 (***) | 0.023 (*) |
| [NiFe] 4a | Evolving: Formate hydrogenlyase | 1 | 0.474 | N/A |
| [NiFe] 4c | Evolving: Carbon monoxide-respiring | 0.123 | 0.685 | 0.163 |
| [NiFe] 4e | Bidirectional: Ech-type | 0.271 | 0.448 | 0.928 |
| [NiFe] 4f | Putative: Formate-coupled | 0.147 | 0.344 | 0.649 |
| [NiFe] 4g | Putative: Ferredoxin-coupled | 0.137 | 0.988 | 0.739 |
| [NiFe] 4h | Uptake: Eha-type | 0.207 | 0.009 (**) | 0.061 |
| [NiFe] 4i | Uptake: Ehb-type | 0.255 | 0.005 (**) | 0.105 |
| **Other hydrogen-metabolizing enzymes** | | **Gene** | **Transcript** | **Ratio** |
| [Fe] | [Fe]-hydrogenase | 0.007 (**) | 0.001 (**) | 0.023 (*) |
| NifH | Nitrogenase | 0.617 | 0.002 (*) | 0.009 (**) |

**Table S9.** Significance testing of differences in hydrogenase gene and transcript abundance between taxonomic orders in the rumen of low and high methane yield sheep. *p* values are shown for differences in abundance of each gene class in the metagenome (column 2), the metatranscriptome (column 3), and the ratio between them (column 4). In all cases, significance was tested using independent two-group Wilcoxon rank-sum tests. No values are given when insufficient reads were detected (N/A).

| **Order** | **Gene** | **Transcript** | **Ratio** |
| --- | --- | --- | --- |
| Bacteroidales | 0.024 (*) | 0.003 (**) | 0.954 |
| Bifidobacteriales | 0.474 | 0.303 | N/A |
| Campylobacterales | 1 | 0.900 | N/A |
| Clostridiales | 0.105 | 0.739 | 0.315 |
| Coriobacteriales | 0.005 (**) | <0.0001 (****) | 0.052 |
| Corynebacteriales | N/A | 1 | N/A |
| Desulfovibrionales | 0.003 (**) | 0.073 | 0.515 |
| Enterobacteriales | 0.736 | 0.482 | N/A |
| Entodiniomorphida | 0.239 | 0.011 (*) | 0.017 (*) |
| Erysipelotrichales | 0.0002 (***) | 0.721 | 0.002 (**) |
| Lactobacillales | <0.0001 (****) | 0.0006 (***) | 0.172 |
| Methanobacteriales | 0.105 | 0.001 (**) | 0.036 (*) |
| Methanomassiliicoccales | 0.089 | 0.045 (*) | 0.839 |
| Neocallimastigales | N/A | 1 | 0.135 |
| Pasteurellales | N/A | 1 | N/A |
| Selenomonadales | 0.684 | 0.046 (*) | 0.315 |
| Spirochaetales | 0.724 | 0.493 | 0.739 |
| Synergistales | 0.695 | 0.922 | 0.424 |

**Table S10.** Significance testing of the differences in gene and transcript abundance of H_2_ uptake pathways in the rumen of low and high methane yield sheep. *p* values are shown for differences in abundance of each gene class in the metagenome (column 3), the metatranscriptome (column 4), and the ratio between them (column 5). In all cases, significance was tested using independent two-group Wilcoxon rank-sum tests.

| **H_2_-metabolizing enzymes** | | **Gene** | **Transcript** | **Ratio** |
| --- | --- | --- | --- | --- |
| NiFe | [NiFe]-hydrogenases (all) | 0.025 (*) | 0.926 | 0.075 |
| FeFe | [FeFe]-hydrogenases (all) | 0.248 | 0.436 | 0.143 |
| DsrA | [Fe]-hydrogenases | 0.007 (**) | 0.002 (**) | 0.110 |
| HydB | [FeFe] A3 diaphorase subunit | 0.015 (*) | 0.247 | 0.631 |
| NifH | Nitrogenase | 0.617 | 0.002 (*) | 0.009 (**) |
| **H_2_ uptake pathways** | | **Gene** | **Transcript** | **Ratio** |
| McrA | Methyl-CoM reductase | 0.105 | <0.0001 (****) | 0.009 (**) |
| AcsB | Acetyl-CoA synthase | 0.118 | <0.0001 (****) | 0.052 |
| AprA | Adenylylsulfate reductase | 0.0003 (***) | 0.217 | 0.025 (*) |
| DsrA | Dissimilatory sulfite reductase | 0.526 | 0.617 | 0.726 |
| AsrA | Alternative sulfite reductase | 0.025 (*) | 0.143 | 0.393 |
| FrdA | Fumarate reductase | 0.401 | 0.002 (**) | 0.215 |
| NarG | Dissimilatory nitrate reductase | 0.839 | 0.223 | 0.149 |
| NapA | Periplasmic nitrate reductase | 0.443 | 0.059 | 0.592 |
| NrfA | Ammonia-forming nitrite reductase | 0.893 | 0.228 | 0.393 |
| DmsA | DMSO / TMAO reductase | 0.726 | 0.0003 (***) | 0.0005 (***) |
| CydA | Cytochrome *bd* oxidase | 0.565 | 0.156 | 0.247 |

**Table S11.** Summary of read mapping results for RNA-seq experiment comparing gene expression of *Ruminococcus albus* 7 and *Wolinella succinogenes* DSM 1740 in pure culture and co-culture.

| **Culture**  **replicate** ^a^ | **Total**  **reads** ^b^ | **Reads after**  **trimming** ^c^ | **Uniquely mapped**  **reads** ^d,e^ | **Unmapped**  **reads** ^f^ |
| --- | --- | --- | --- | --- |
| *R. albus* 1 | 31.2 M | 31.1 M | 77.1% | 21.2% |
| *R. albus* 2 | 31.4 M | 31.2 M | 80.0% | 18.7% |
| *W. succinogenes* 1 | 29.3 M | 29.1 M | 75.0% | 21.6% |
| *W. succinogenes* 2 | 25.5 M | 25.4 M | 74.2% | 22.4% |
| Co-culture 1 | 28.4 M | 30.7 M | 35.0% (Ra),  42.0% (Ws) | 23.0% |
| Co-culture 2 | 33.3 M | 33.1 M | 37.2% (Ra),  38.5% (Ws) | 24.3% |

^a^ Cultures of *R. albus* and *W. succinogenes* were grown in modified Balch medium until mid-log phase before RNA was extracted from two biological replicates and sequenced.

^b^ Sequenced reads were quantified for each sample. M = million.

^c^ Reads were trimmed for quality and quantified. M = million.

^d^ Reads were mapped to either the *R. albus* genome (Ra) or the *W. succinogenes* (Ws) genome.

^e^ The number of reads mapped to one segment of the genome. Reads were only mapped if the fraction of the read that aligned with the reference genome was greater than 0.9 and if the read matched other regions of the reference genome at less than 10 nucleotide positions.

^f^ The number of reads not mapped to the genomes, either because they were too short or did not map to the coding region of the genome.

**Figure S1. Distribution of hydrogenase subgroups in rumen microorganisms.** Results are shown based on screens of the 501 genomes of cultured bacteria, archaea, and eukaryotes (**Table S2)**. The hydrogenases were classified into phylogenetically and functionally distinct subgroups based on the scheme of HydDB. Partial hydrogenase sequences were also retrieved and classified from four rumen ciliates and two rumen fungi. **Table S1** lists the FASTA sequences of the retrieved reads and **Table S2** shows the distribution of these enzymes.

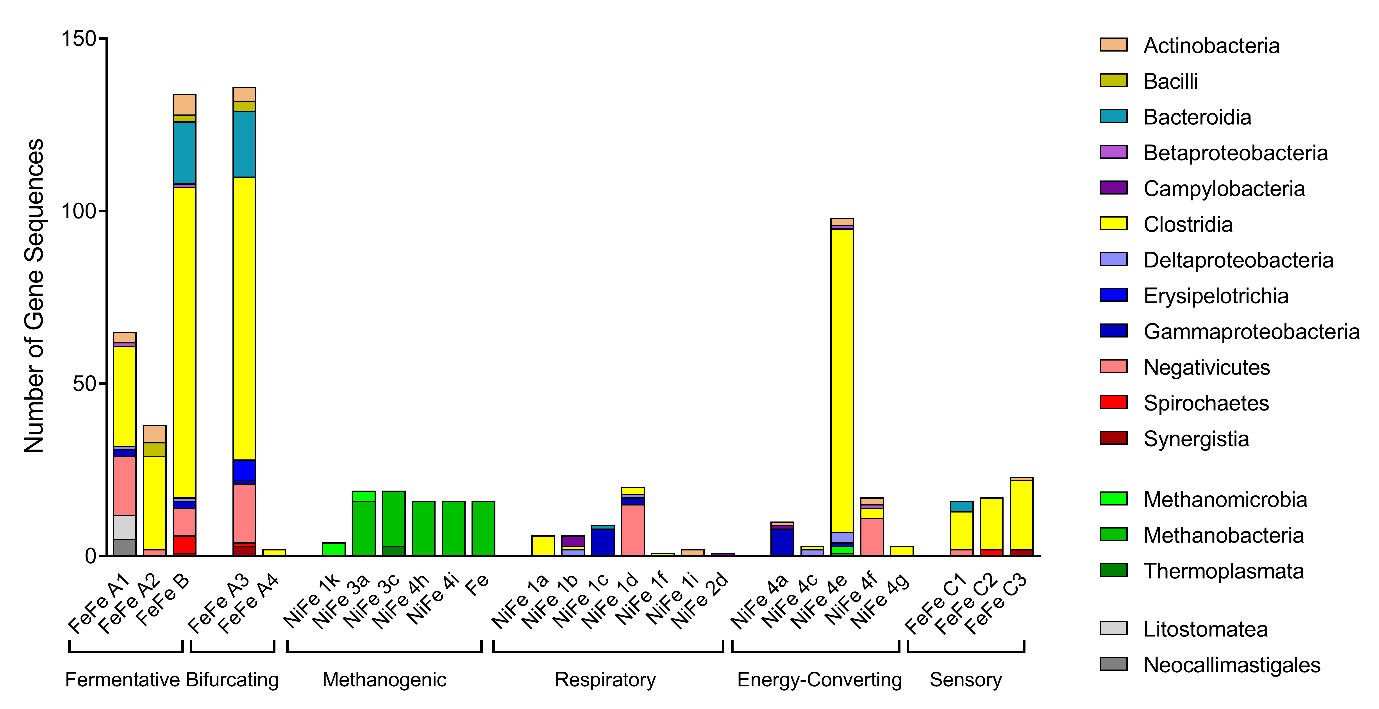

**Figure S2. Comparison of abundance of different hydrogenases, divided by hydrogenase subgroup, in metagenome and metatranscriptome datasets from the rumen of low and high methane yield sheep.** The normalized count of metagenome reads (a), normalized count of metatranscriptome reads (b), and ratio of metatranscriptome to metagenome reads (c) are shown. Results are shown for ten paired metagenome and metatranscriptome datasets each from low methane yield sheep (orange) and high methane yield sheep (blue) that were randomly subsampled at five million reads. Each boxplot shows the ten datapoints and their range, mean, and quartiles. Significance was tested using independent two-group Wilcoxon rank-sum tests (* *p* < 0.05; ** *p* < 0.01; *** *p* < 0.001; **** *p* < 0.0001; full *p* values in **Table S8**). Note that the metatranscriptome data is also shown in **Figure 3a**, but is shown again here to facilitate comparison of metagenome and metatranscriptome data.

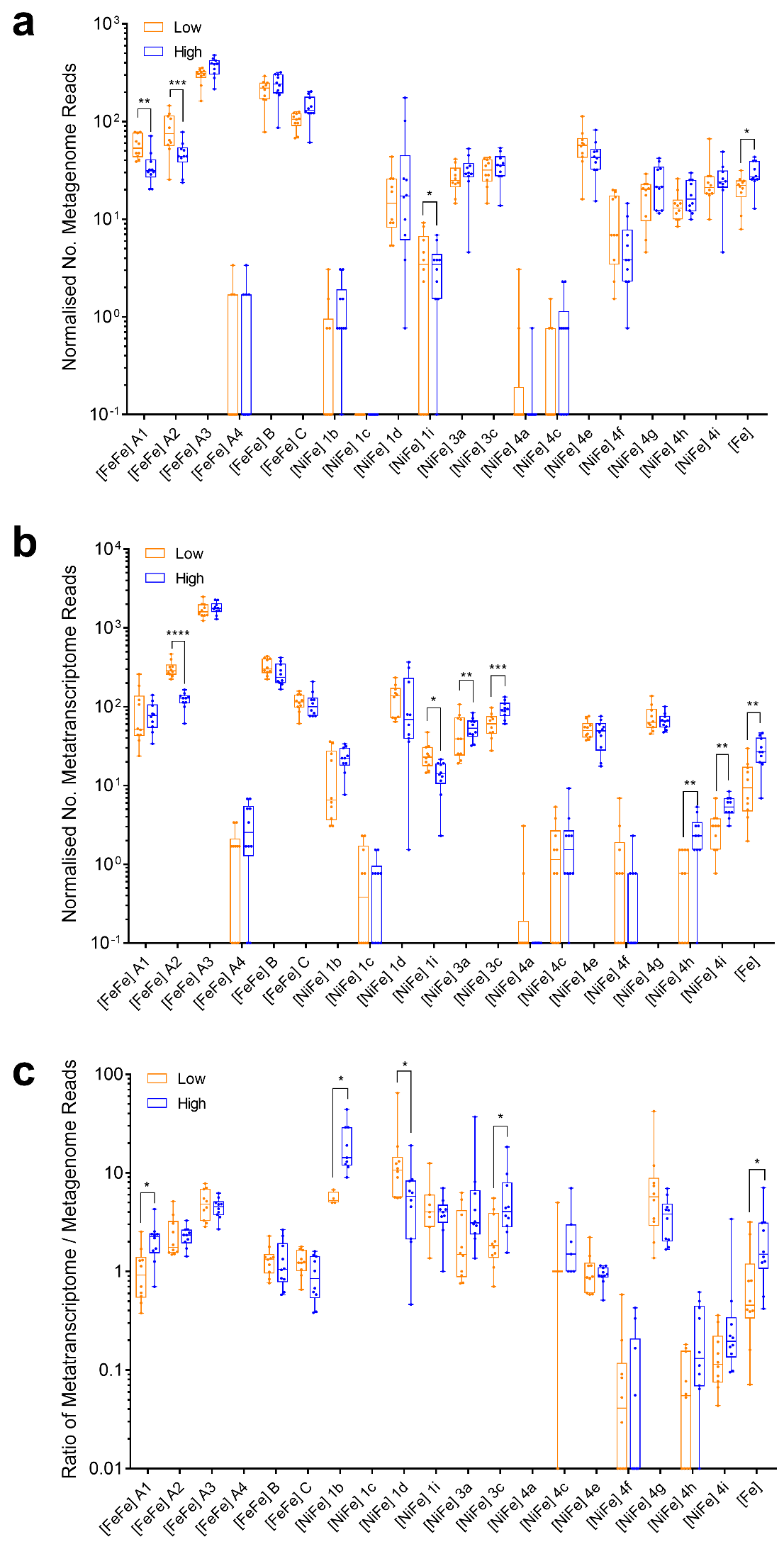

**Figure S3. Comparison of abundance of different hydrogenases, as divided by predicted taxonomic affiliation, in metagenome and metatranscriptome datasets from the rumen of low and high methane yield sheep.** The normalized count of metagenome reads (a), normalized count of metatranscriptome reads (b), and ratio of metatranscriptome to metagenome reads (c) are shown. Results are shown for ten paired metagenome and metatranscriptome datasets each from low methane yield sheep (orange) and high methane yield sheep (blue) that were randomly subsampled at five million reads. Each boxplot shows the ten datapoints and their range, mean, and quartiles. Significance was tested using independent two-group Wilcoxon rank-sum tests (* *p* < 0.05; ** *p* < 0.01; *** *p* < 0.001; **** *p* < 0.0001; full *p* values in **Table S9**). Note that the metatranscriptome data is also shown in **Figure 3b**, but is shown again here to facilitate comparison of metagenome and metatranscriptome data.

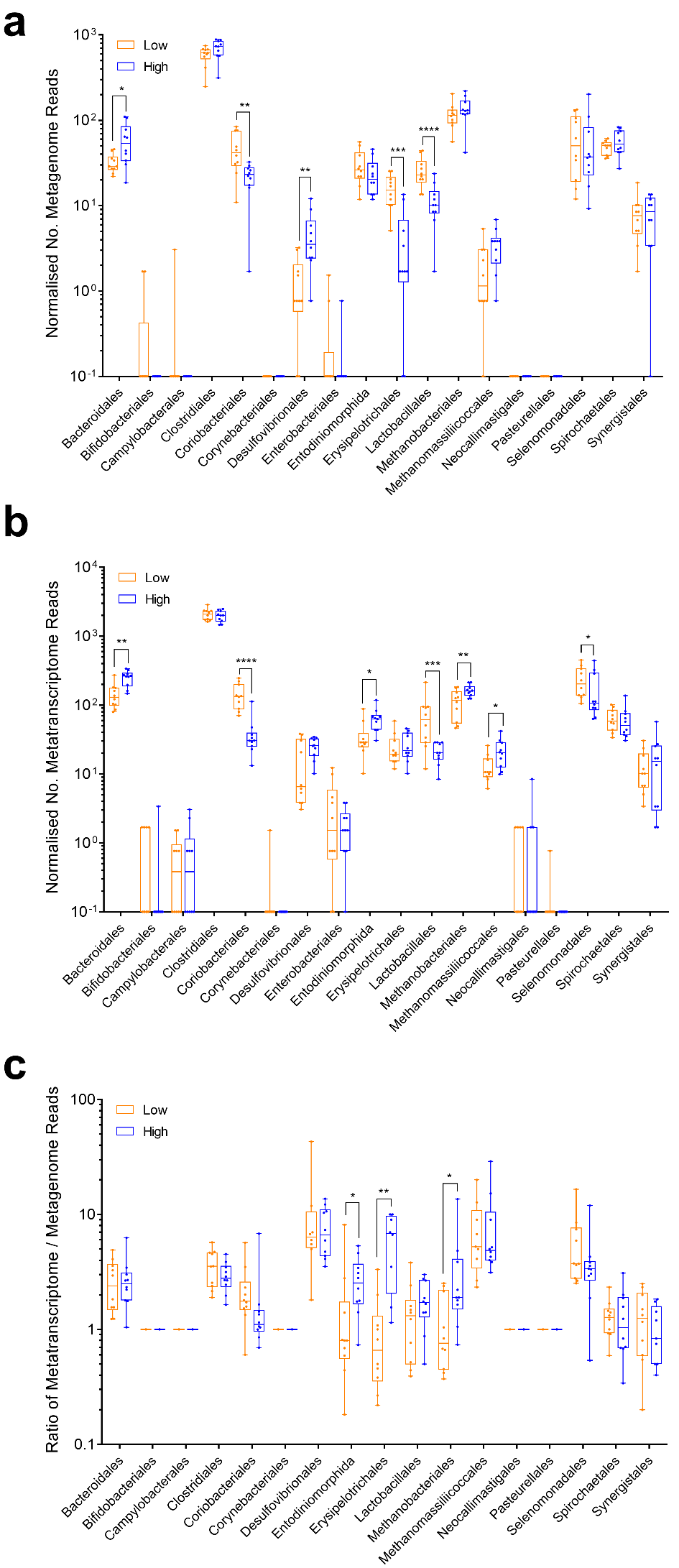

**Figure S4. Comparison of abundance of different H_2_ uptake pathways in metagenome and metatranscriptome datasets from the rumen of low and high methane yield sheep.** The normalized count of metagenome reads (a), normalized count of metatranscriptome reads (b), and ratio of metatranscriptome to metagenome reads (c) are shown. The boxplots show the number of transcripts encoding the catalytic subunits of the [NiFe]-hydrogenases (NiFe), [FeFe]-hydrogenases (FeFe), [Fe]-hydrogenases (Fe), hydrogenase-associated diaphorase (HydB), nitrogenases (NifH), methyl-CoM reductases (McrA), acetyl-CoA synthases (AcsB), adenylylsulfate reductases (AsrA), dissimilatory sulfite reductases (DsrA), anaerobic sulfite reductases (AsrA), fumarate reductases (FrdA), dissimilatory nitrate reductases (NarG), periplasmic nitrate reductase (NapA), ammonia-forming nitrite reductases (NrfA), DMSO / TMAO reductases (DmsA), and cytochrome *bd* oxidases (CydA). For fumarate reductase and cytochrome *bd* oxidase, the numerous reads from non-hydrogenotrophic organisms (e.g. Bacteroidetes) were excluded. Results are shown for ten paired metagenome and metatranscriptome datasets each from low methane yield sheep (orange) and high methane yield sheep (blue) that were randomly subsampled at five million reads. Each boxplot shows the ten datapoints and their range, mean, and quartiles. Significance was tested using independent two-group Wilcoxon rank-sum tests (* *p* < 0.05; ** *p* < 0.01; *** *p* < 0.001; **** *p* < 0.0001; full *p* values in **Table S10**). Note that the metatranscriptome data is also shown in **Figure 3c**, but is shown again here to facilitate comparison of metagenome and metatranscriptome data.

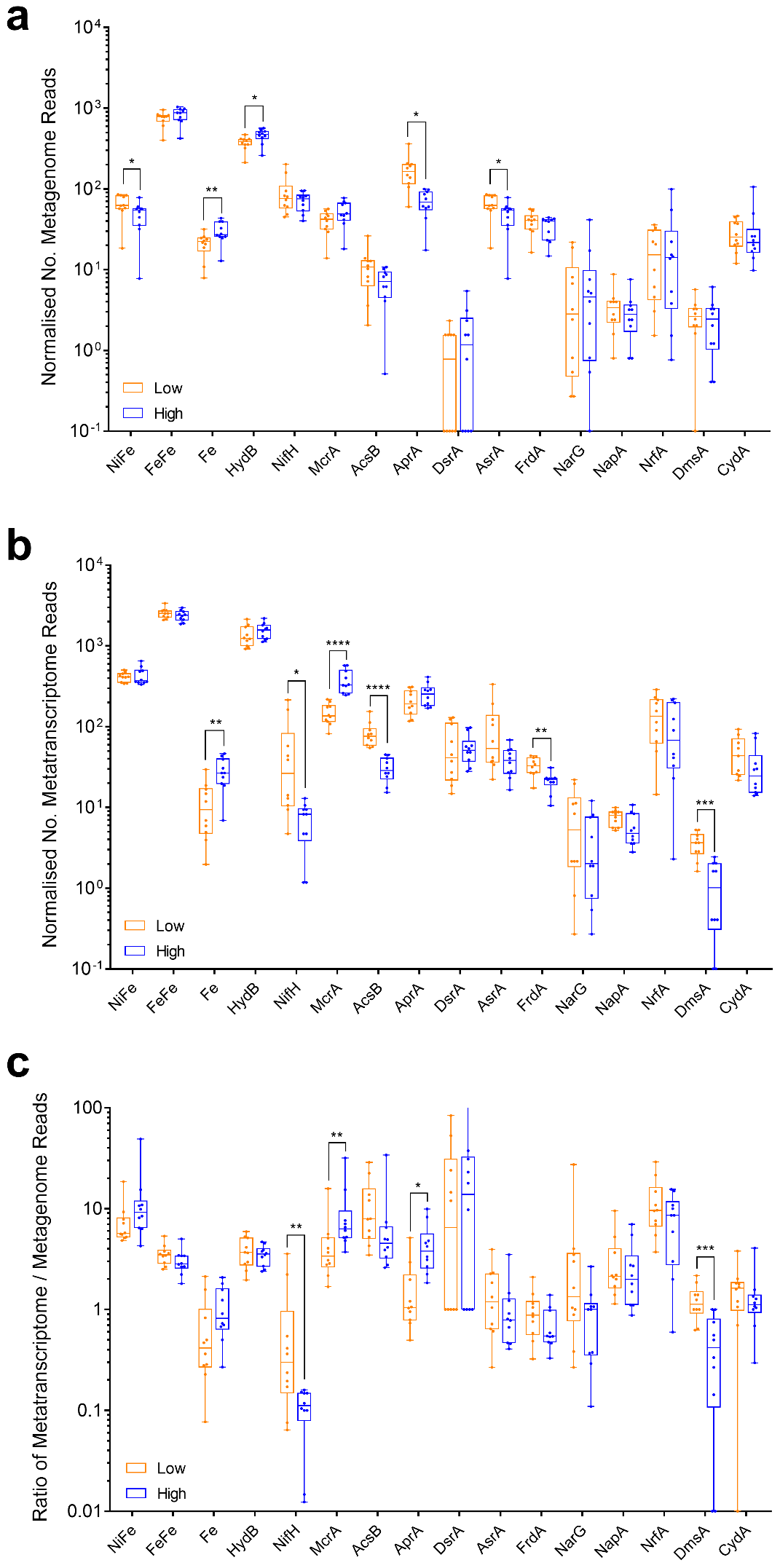

**Figure S5.** **Comparison of normalized expression levels of the most highly abundant hydrogenases from the rumen of low and high methane yield sheep.** (a) Normalized metatranscriptome read count of 20 predicted H_2_-evolving hydrogenases. (b) Normalized metatranscriptome read count of 20 predicted H_2_-uptake and putative hydrogenases. The group 1b, 1d, 1i, 3a, 3c [NiFe]-hydrogenases and [Fe]-hydrogenase mediate H_2_ uptake. In acetogenic organisms, the group A3 [FeFe]-hydrogenases can also mediate H_2_ uptake. The directionality and role of the group 4g [NiFe]-hydrogenases and group A2 [FeFe]-hydrogenases have not been experimentally determined. Each boxplot shows the ten datapoints and their range, mean, and quartiles. Significance was tested using independent two-group Wilcoxon rank-sum tests (* *p* < 0.05; ** *p* < 0.01; *** *p* < 0.001; **** *p* < 0.0001). A full list of metagenome and metatranscriptome hits is provided in **Table S5**.

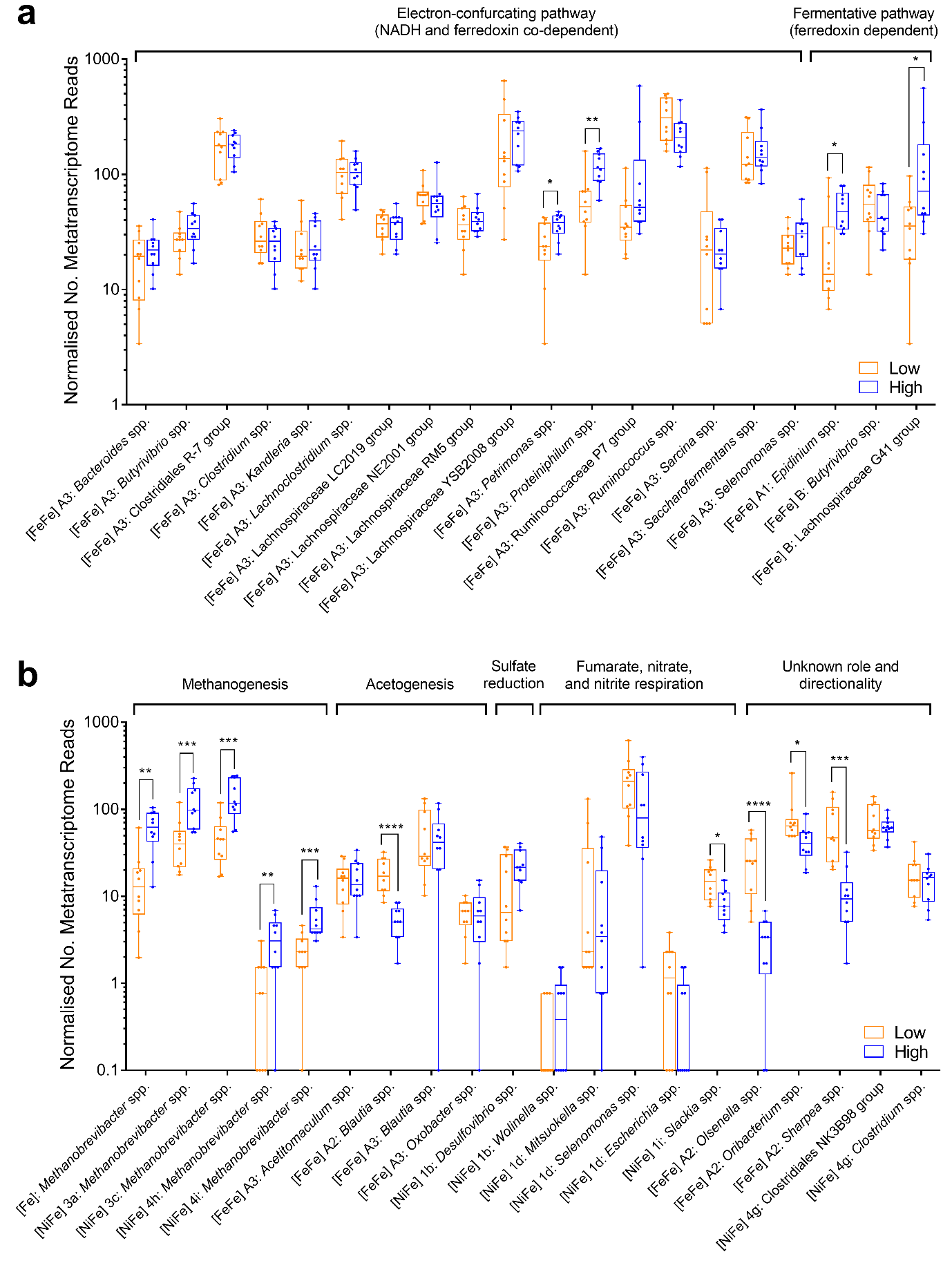

**Figure S6. Comparison of taxonomic assignment of hydrogenase reads from the metatranscriptomes of low and high methane yield sheep at different sequence identity cutoffs.** The predicted taxonomic affiliation of the hydrogenase reads is shown by genera at sequence identity cutoffs of 65% (a) and 85% (b). Genera with greater than 2.5% relative abundance are shown.

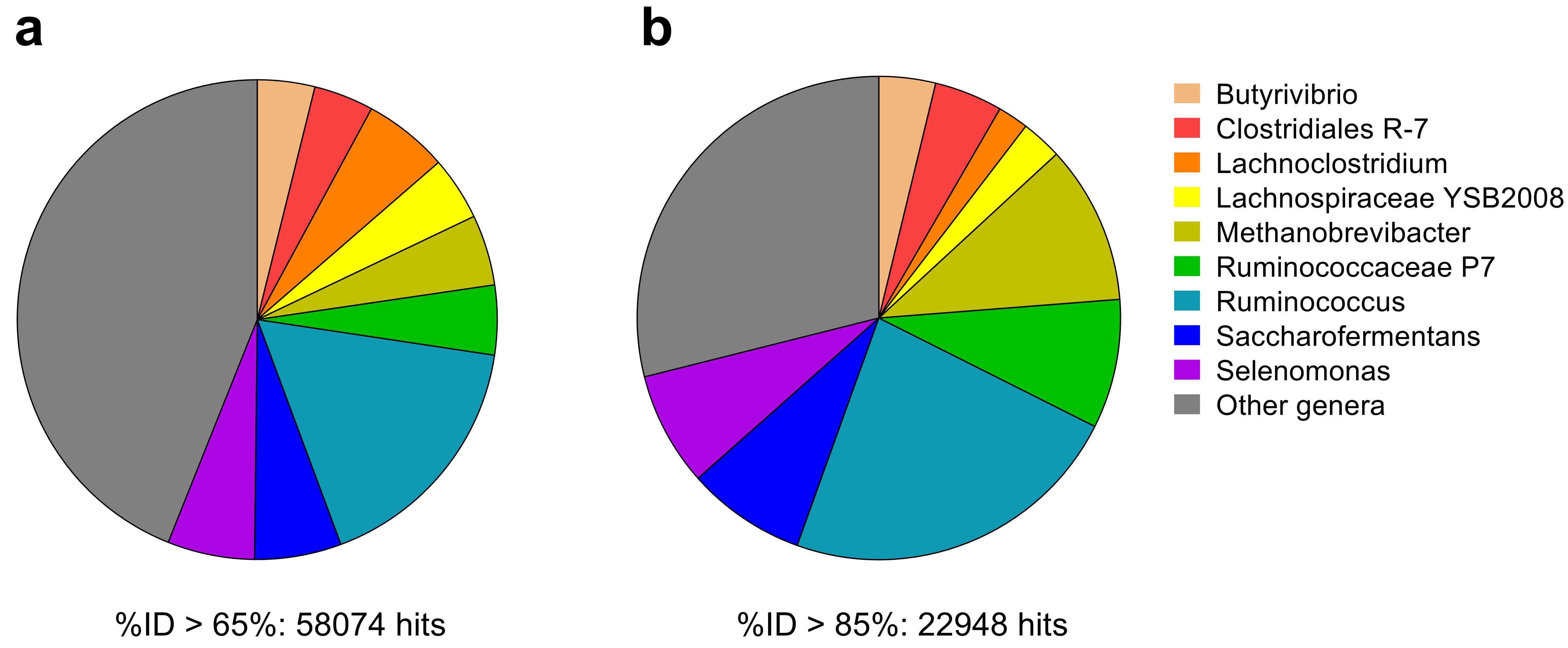

**Figure S7.** **Taxonomic distribution of metatranscriptomic reads encoding hydrogenases and potential H_2_ uptake pathways from the rumen of low and high methane yield sheep.** Reads assigned to orders lacking hydrogenases are shown in grey.

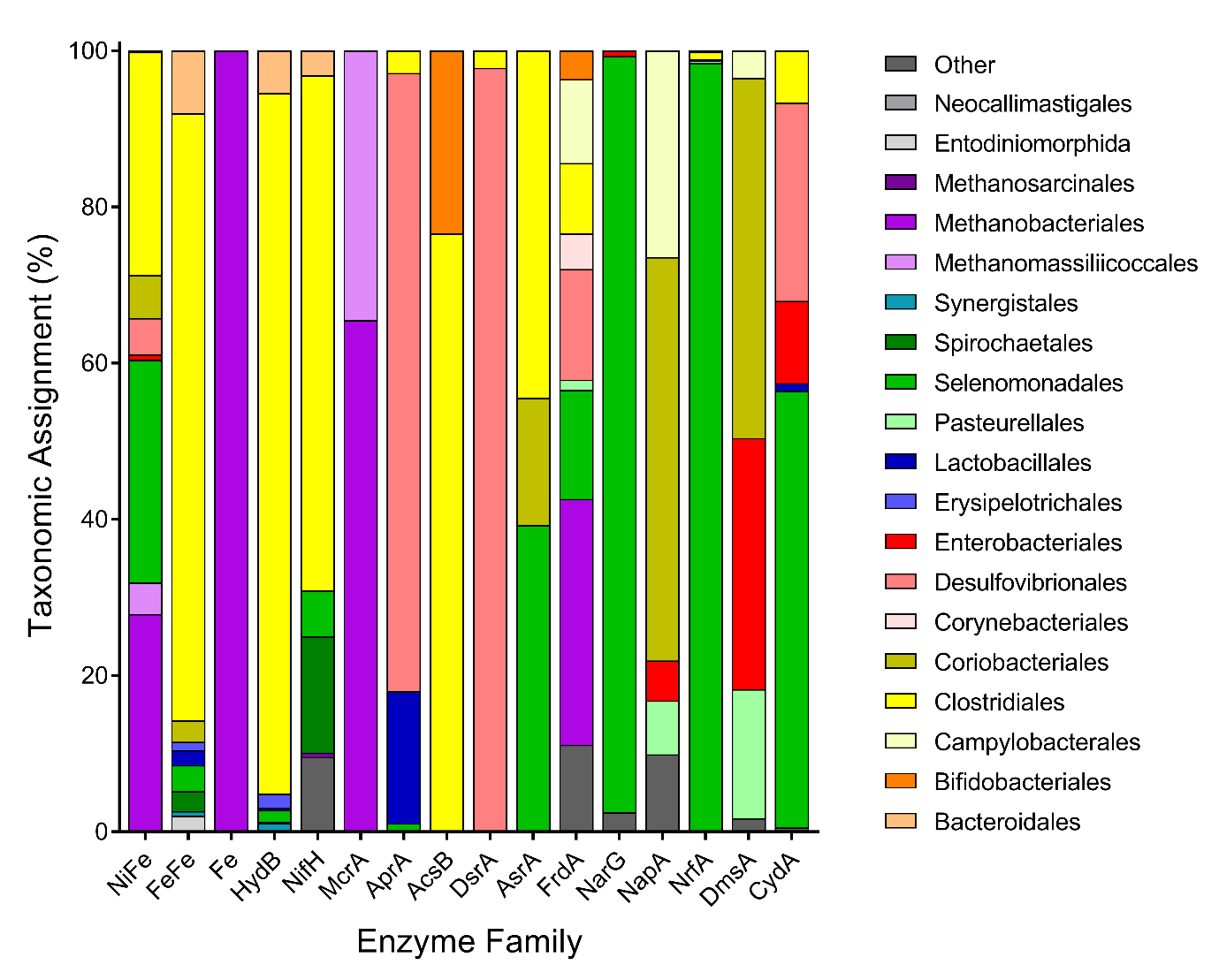

**Figure S8. Changes in metabolite concentrations during growth of *Ruminococcus albus* and *Wolinella succinogenes* in pure culture and co-culture.** Results are shown for *Ruminococcus albus* 7 in pure culture (a), *Wolinella succinogenes* DSM 1740 in pure culture (b), and a co-culture of both strains (c). Data are reported as means ± standard deviation from three biological replicates. No formate was detected under any of the three culture conditions.

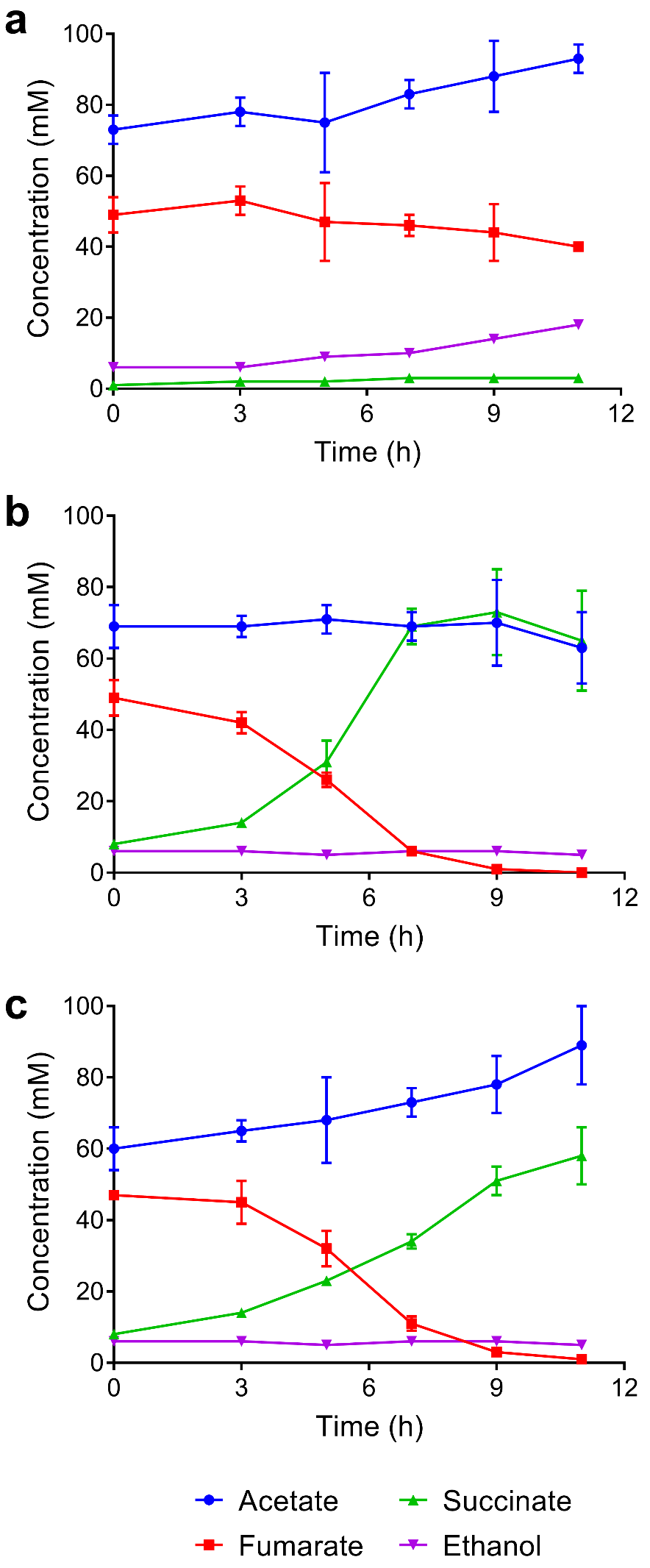
